# Context-specific functions of Notch in *Drosophila* blood cell progenitors

**DOI:** 10.1101/682658

**Authors:** DM Blanco-Obregon, MJ Katz, L Durrieu, L Gándara, P Wappner

**Author notes:** Contributed equally to this work (DM Blanco-Obregon, MJ Katz). **Summary statement** Notch signaling regulates differently distinct populations of blood cell progenitors of the *Drosophila* larval hematopoietic organ.

## Abstract

*Drosophila* Larval hematopoiesis takes place at the lymph gland, where myeloid-like progenitors differentiate into Plasmatocytes and Crystal Cells, under regulation of conserved signaling pathways. It has been established that the Notch pathway plays a specific role in Crystal Cell differentiation and maintenance. In mammalian hematopoiesis, the Notch pathway has been proposed to fulfill broader functions, including Hematopoietic Stem Cell maintenance and cell fate decision in progenitors. In this work we describe different roles that Notch plays in the lymph gland. We show that Notch, activated by its ligand Serrate, expressed at the Posterior Signaling Center, is required to restrain Core Progenitor differentiation. We define a novel population of blood cell progenitors that we name Distal Progenitors, where Notch, activated by Serrate expressed in Lineage Specifying Cells at the Medullary Zone/Cortical Zone boundary, regulates a binary decision between Plasmatocyte and Crystal Cell fates. Thus, Notch plays context-specific functions in different blood cell progenitor populations of the *Drosophila* lymph gland.

**Graphical Abstract:** 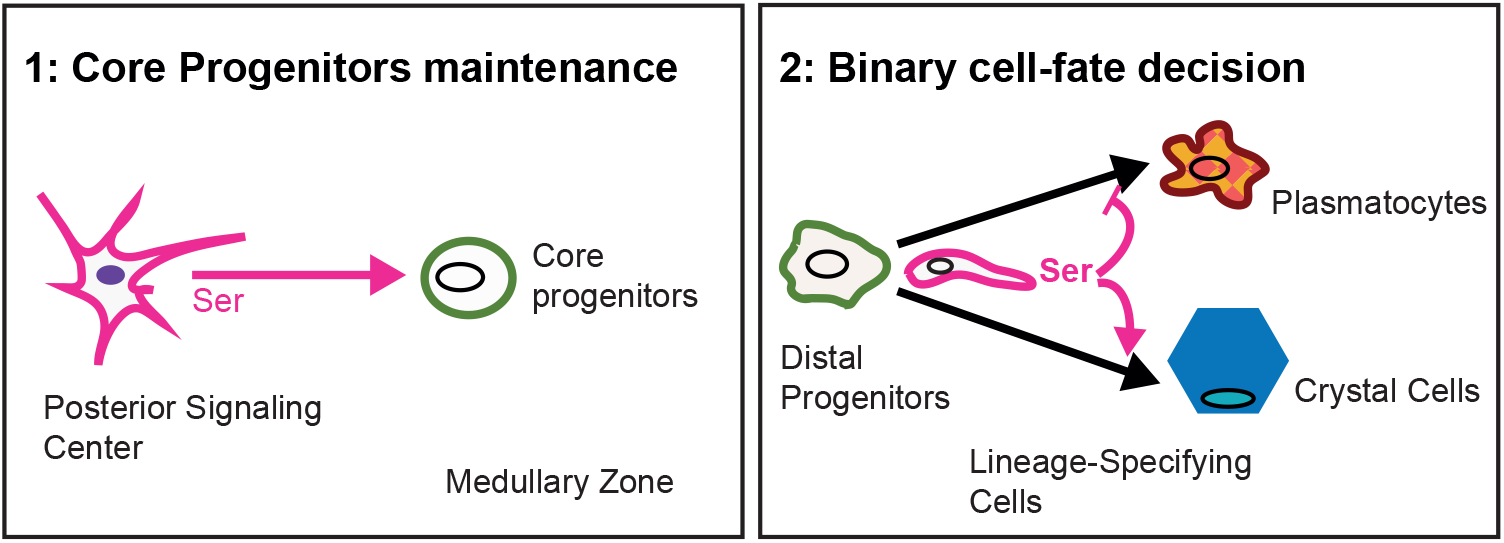

## Introduction

Hematopoiesis in *Drosophila* larvae takes place predominantly at the Lymph Gland, which is composed of two primary lobes symmetrically localized at both sides of the dorsal vessel, and smaller posterior lobes that follow a similar array (Jung et al., 2005) At the primary lobes, blood cell progenitors differentiate into myeloid-like lineages through evolutionary conserved mechanisms, making the lymph gland an attractive model to explore the pathways involved in normal and oncogenic hematopoiesis (Banerjee et al., 2019; Crozatier and Vincent, 2011; Evans et al., 2007). At the 3^rd^ larval instar, blood cell progenitors are compactly arranged in an internal region of the primary lobe, called Medullary Zone (MZ), characterized by the expression of the JAK/STAT receptor *domeless* (*dome*) (**Fig. 1A**) (Evans et al., 2014; Jung et al., 2005; Krzemien et al., 2007). Maturing hemocytes are found in the peripheric region of the lobe, called Cortical Zone (CZ), which can be identified by the expression of the Von Willebrand-like factor *Hemolectin* (*Hml*) (**Fig. 1A, B**) (Jung et al., 2005). An Intermediate Progenitor (IP) population has been described at the MZ/CZ boundary, defined by the expression of both *dome* and *Hml* (**Fig. 1A, B**), but its physiological relevance remains poorly defined (Banerjee et al., 2019; Krzemien et al., 2010a). Two types of differentiated populations of hemocytes occur at the CZ: Plasmatocytes (PLs) and Crystal Cells (CCs) (**Fig. 1A**), whereas a third cell type, the Lamellocytes, differentiate only following specific immune challenges. PLs, which constitute the bulk (95%) of mature hemocytes, are macrophages that retain *Hml* expression, while they also express the receptors Nimrod C1 (or P1-antigen) and eater, both of them important for recognition of bacteria (Kocks et al., 2005; Kurucz et al., 2007a; Kurucz et al., 2007b). The CCs, named after their cytoplasmic inclusions of Prophenoloxydase (ProPO), mediate melanization of pathogens and wounds, and constitute 5% of the total number of mature hemocytes. Mature CCs are thus characterized by the expression of ProPO, and no longer express the *Hml* marker, characteristic of other CZ cells (Evans et al., 2014; Rizki et al., 1980). A population of CC Progenitors can also be identified at the CZ by the expression of the RUNX transcription factor lozenge (Lz), homolog of the human AML1/Runx1 protein, frequently altered in Acute Myeloid Leukemias (Jung et al., 2005; Lebestky, 2000). Another remarkable lymph gland region, termed Posterior Signaling Center (PSC), occurs at the posterior tip of each primary lobe (**Fig. 1B**). It has been reported to function as the niche that maintains the progenitor population of the MZ in an undifferentiated state (Mandal et al., 2007; Mondal et al., 2011; Pennetier et al., 2012). PSC cells express the homeobox protein Antennapedia (Antp) (Mandal et al., 2007), the signaling molecule hedgehog (Hh) (Mandal et al., 2007), the Notch ligand Serrate (Ser) (Lebestky et al., 2003), and the gene *collier* (*col*), ortholog of the mammalian Early B-cell Factor (EBF) (Crozatier et al., 2004; Krzemien et al., 2007). Hh expressed at the PSC targets the MZ and is required to restrain progenitor differentiation (Mandal et al., 2007). Recently the notion that the PSC functions as a hematopoietic niche has been challenged, as genetic ablation of this structure did not alter progenitor maintenance nor steady-state blood cell differentiation (Baldeosingh et al., 2018; Benmimoun et al., 2015).

**Figure 1.**
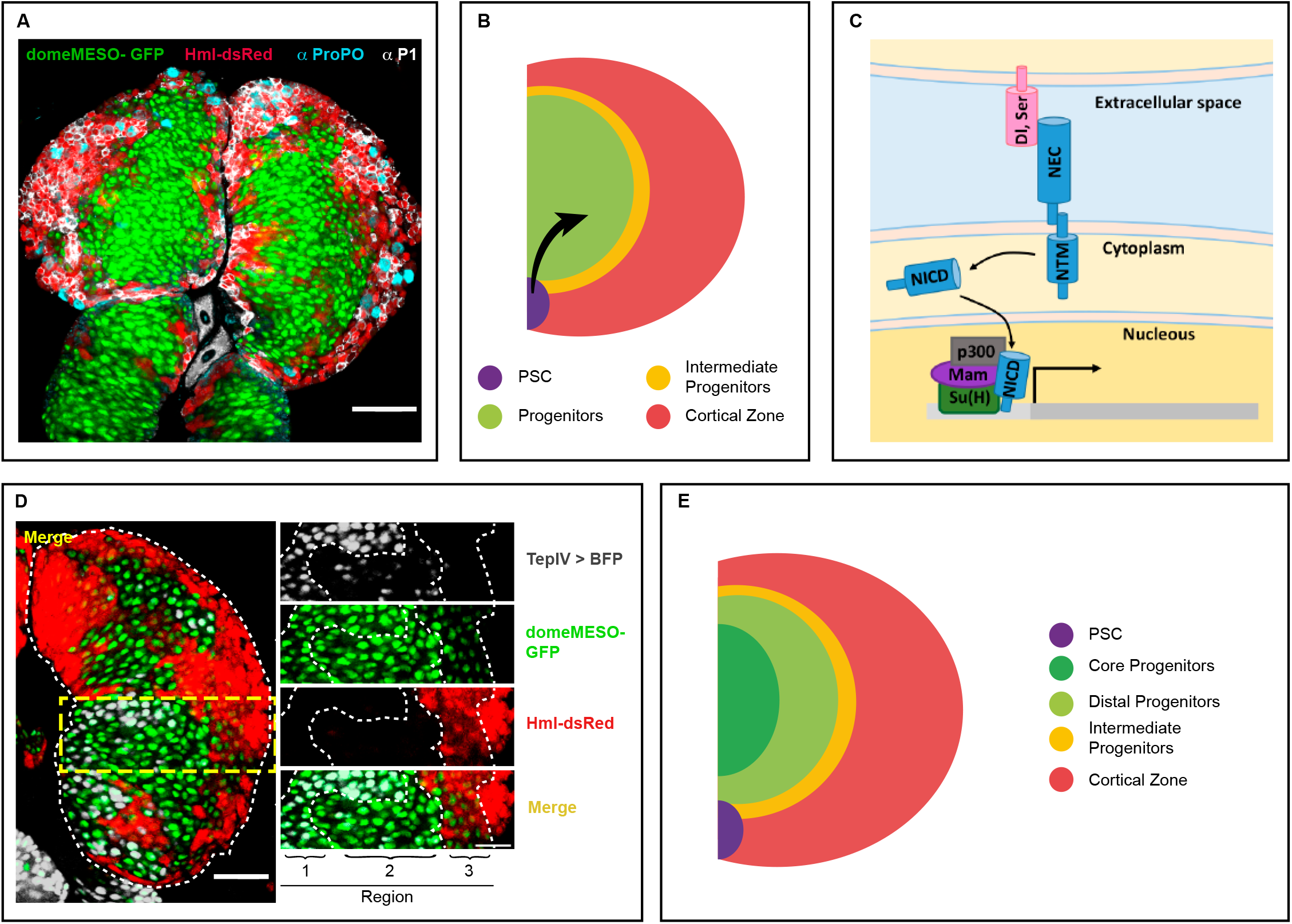
Redefinition of the lymph gland progenitor subpopulations. **(A) Cell populations of the lymph gland.** Confocal image of a lymph gland from a 3^rd^ instar wandering larva. Blood cell progenitors (green; *domeMESO-*GFP); differentiating cells (red; *Hml*-dsRed); Crystal Cells (cyan, anti-ProPO staining), and Plasmatocytes (white, anti-P1 staining). Scale bar, 50 μm. **(B) Current view on the cell populations of the lymph gland.** The scheme illustrates a cross section of a wandering 3^rd^ instar larva lymph gland lobe (anterior is up and center is to the left), depicting the described populations in different colors. Purple: Posterior Signaling Center (PSC); green: Blood Cell Progenitors; yellow: Intermediate Progenitors; pink: Cortical Zone where differentiating cells, Plasmatocytes and Crystal Cells are localized. The black arrow indicates progenitor maintenance signals. **(C) Simplified diagram of the Notch pathway.** Representation of the canonical Notch Pathway in *Drosophila*. Dl: Delta, Ser: Serrate. NED: Notch Extracellular Domain. NTM: Notch Transmembrane Fragment. NICD: Notch Intracellular Domain, Su(H): Supressor of Hairless, Mam: Mastermind. **(D) Three populations of blood cell progenitors occur at the lymph gland.** Single-plane confocal images of a primary lobe from wandering third instar larvae, where three regions can be defined based on differential marker expression. Region 1: Cells expressing both *TepIV* > BFP (white) and *domeMESO*-GFP (green). Region 2: Cells that express *domeMESO*-GFP but not *TepIV* > BFP nor *Hml*-dsRed (red). Region 3: Cells in which *domeMESO-*GFP and *Hml*-dsRed signals coexist. The dashed line indicates the outline of the primary lobe or the limits of progenitors subpopulations in the inset. Scale bar, 50 μm. **(E) Schematic representation of the redefined progenitor populations.** A wandering 3^rd^ instar wild-type lymph gland lobe is represented (cross section, anterior is up and center is to the left). Progenitors encompass three subpopulations: 1) Core Progenitors, positive for *domeMESO* and *TepIV* (dark green); 2) Distal Progenitors, positive for *domeMESO* (light green) but negative for *TepIV* and *Hml*; and 3) Intermediate Progenitors, which are positive for both *domeMESO* and *Hml* (yellow). Purple: Posterior Signaling Center (PSC); pink: Cortical Zone.

Notch is a conserved signaling pathway utilized repeatedly during development of all metazoa. It is typically involved in cell differentiation, binary cell fate decisions, cell proliferation and cell survival (Bray, 2016). The Notch receptor, which gives the name to the pathway, can be activated by its transmembrane ligands Serrate (Ser) or Delta (Dl) expressed in adjacent cells (**Fig. 1C**). Once this interaction takes place, Notch undergoes two consecutive cleavage events, resulting in the release of the Notch Intracellular Domain (NICD), which migrates into the nucleus and binds the transcription factor Suppressor of Hairless (Su(H)). The complex formed by NICD and Su(H) recruits transcriptional co-activators, thereby inducing transcription of Notch target genes. This canonical Notch pathway has been reported to operate in *Hml*-positive cells of the CZ to specify the CC fate (Duvic et al.; Lebestky et al., 2003; Mukherjee et al., 2011). The source of Notch ligand in this context is Ser expressed in Lineage Specifying Cells (LSCs) localized at the MZ/CZ boundary (Ferguson and Martinez-Agosto, 2014; Lebestky et al., 2003). Non-canonical ligand-independent Notch signaling is afterwards required for CC maturation and survival (Mukherjee et al., 2011).

In mammalian hematopoiesis, Notch pathway functions have been explored quite extensively, although with contrasting results. Notch receptors (Notch 1-4) are expressed in Hematopoietic Stem Cells (HSCs), hematopoietic progenitors and mature blood cells, suggesting that Notch is required at multiple stages of the differentiation cascade (Liu et al., 2010). Notch functions in mammalian HSCs are controversial (Lampreia et al., 2017). Results in mouse models suggest that it is dispensable for their maintenance (Maillard et al., 2008; Mancini et al., 2005), however *in vitro* conflicting evidence favoring either a function of Notch in HSC differentiation (Schroeder and Just, 2000; Schroeder et al., 2003), or a requirement for HSC maintenance have been reported (Kumano et al., 2001; Varnum-Finney et al., 2000; Vercauteren and Sutherland, 2004). It is however well established that in lymphoid progenitors Notch promotes differentiation and proliferation of T lymphocytes at the expense of B lymphocytes (Han et al., 2002; Radtke et al., 1999). Given the involvement of Notch in mammalian hematopoiesis, it is not surprising that alterations of this pathway are associated with several types of leukemia (Kushwah et al., 2014).

Because of the diverse, yet unclear functions of Notch in mammalian hematopoiesis, and given the conservation that occurs between the mechanisms controlling fly and mammalian blood cell development, we sought to explore the functions that the Notch pathway plays in *Drosophila* blood cell progenitors. We show here that Notch has distinct functions in two different populations of hemocyte progenitors: 1) In the recently described “Core progenitors” (Oyallon et al., 2016) Notch is required for maintenance of an undifferentiated state, a function that depends on Ser expressed at the PSC; and 2) In Distal Progenitors, a cell population defined in this paper, Notch controls a binary decision towards a PL or a CC fate. This binary Notch dependent choice depends on Ser expressed in LSCs (Ferguson and Martinez-Agosto, 2014) at the MZ/CZ boundary. Thus, Notch plays context-specific functions in two different cell progenitor populations during *Drosophila* hematopoiesis.

## Results

### Redefining progenitor cell populations of the Medullary Zone

Before analyzing Notch functions in *Drosophila* blood cell progenitors, we sought to define precisely the different populations of progenitors that occur at the MZ. Cells of the MZ of wandering 3^rd^ instar larvae lymph glands express *domeMESO* and include a subpopulation of internal cells, the Core Progenitors, characterized by the expression of *TepIV* (**Fig. S1)**(Oyallon et al., 2016). It is unclear in the literature whether all *domeMESO-* expressing cells that are negative for *TepIV* coexpress *Hml*, and can be considered Intermediate Progenitors. We found that this is not the case: In larvae that coexpress the Blue Fluorescent Protein (BFP) under a *TepIV*-Gal4 driver (*TepIV* > BFP), and GFP directly driven by a d*omeMESO* promoter (*domeMESO*-GFP), along with dsRed controlled directly by an *Hml* promoter (*Hml*-dsRed), three distinct cell populations can be recognized: 1) Core Progenitors positive for both *TepIV* > BFP and *domeMESO*-GFP (**Fig. 1D**, region 1 and **Fig. S1**); 2) Cells positive only for *domeMESO*-GFP (**Fig. 1D**, region 2 and **Fig. S1**); and 3) Intermediate Progenitors in which *domeMESO*-GFP and *Hml*-dsRed signals overlap (**Fig. 1D**, region 3 and **Fig. S1**). Thus, an uncharacterized population of *domeMESO* progenitors which do not express *TepIV* or *Hml* occurs in the lymph gland. We henceforth propose the name “Distal Progenitors” for this particular population, in reference to their distal location from the Core Progenitors (**Fig 1E**).

In conclusion, three distinct populations of hemocyte progenitors occur at the MZ of 3^rd^ instar larvae lymph glands: 1) Core Progenitors, which co-express *TepIV* and *domeMESO*; 2) Distal Progenitors that are positive for *domeMESO*, and negative for *TepIV* and *Hml*; and 3) Intermediate Progenitors, which co-express *domeMESO* and *Hml* (**Fig. 1E**).

### The Notch pathway is required for Core Progenitor maintenance

Notch expression is widespread throughout the lymph gland, suggesting that this pathway might operate in various cell types (**Fig. 2A**). We initially analyzed Notch function in Core Progenitors by expressing a *Notch* RNAi with *TepIV*-Gal4 (**Fig 2B; Fig. S2A**), and observed a clear reduction of Core Progenitors (**Fig. 2B**), while both Plasmatocytes (PLs) and Crystal Cells (CCs), increased significantly (**Fig. 2B; Fig. S2B). Silencing of N** produced no detectable alteration of progenitor cell size **(Fig. S2C)** or effects on the PSC **(Fig. S2D**). Consistent with this, silencing of another component of the canonical Notch pathway, the transcription factor Suppressor of Hairless (Su(H)), also provoked a reduction of Core Progenitors, while PLs and CCs increased significantly (**Fig. 2C, Fig. S2E**). Altogether, these results suggest that the Notch pathway is required cell-autonomously for maintenance of Core Progenitors in an undifferentiated state (**Fig. 2F)**.

**Figure 2.**
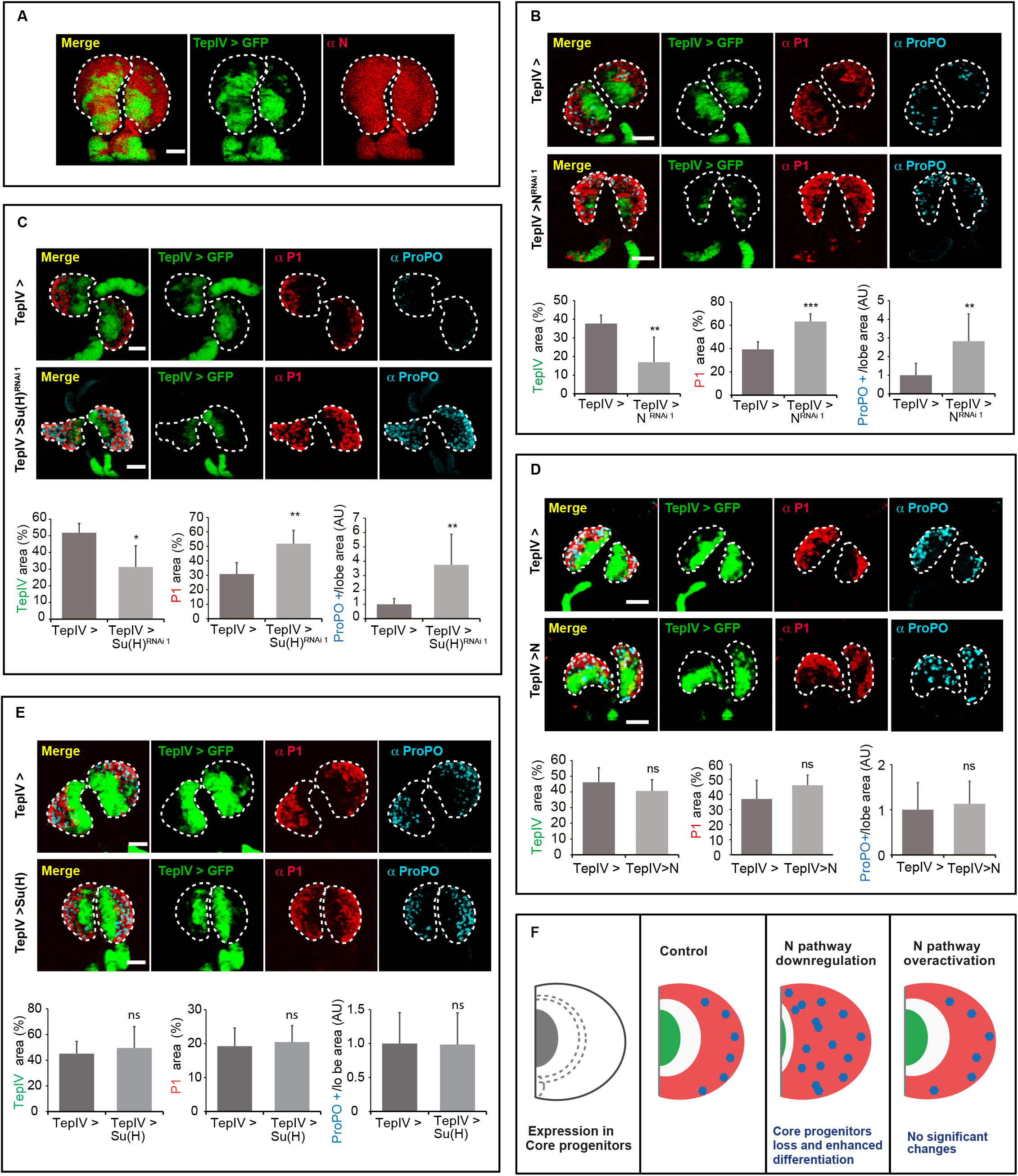
The Notch pathway is required for Core Progenitor maintenance. **(A) *Notch* is expressed throughout the lymph gland.** Staining with an anti-Notch antibody (red) in lymph glands, where the Core Progenitors are marked with *TepIV*-Gal4/*UAS-GFP* (green). Single-plane confocal images of primary lobes from wandering third instar larvae are shown. The dashed line marks the outline of the lymph gland lobes. Scale bars, 50 μm. **(B) Notch is required for Core Progenitor maintenance.** *Notch RNAi* (*N^RNAi^ ^1^*) expression in Core Progenitors, driven by *TepIV*-Gal4, provokes loss of Core Progenitors (green: *TepIV* > GFP), along with increased Plasmatocytes (red: P1 staining) and Crystal Cells (cyan: ProPO staining). Compare control lymph glands in upper panels with lymph glands expressing *N*^RNAi^ ^1^ in middle panels. Whole Z-projections of confocal images of primary lobes from wandering third instar larvae are depicted. The dashed line marks the outline of the two lymph gland lobes. Scale bars, 60 μm. Graphs in lower panels show quantification of the indicated markers, normalized to the area of the primary lobe (**p < 0.01, ***p < 0.001). Error bars represent SD. *TepIV* >, n = 8; *TepIV* > *N*^RNAi 1^, n = 8. **(C) Supressor of Hairless (Su(H)) is required for Core Progenitor maintenance.** *Su(H)*^RNAi1^ expression with *TepIV*-Gal4 caused reduction of Core Progenitors (green: *TepIV* > GFP), and simultaneous increase of Plasmatocytes (red: P1 staining) and Crystal Cells (cyan: ProPO staining). Compare control lymph glands in upper panels with lymph glands expressing *Su(H)*^RNAi1^ in middle panels. Whole Z-projections of confocal images of primary lobes from wandering third instar larvae are represented. The dashed line indicates the outline of the two primary lobes. Scale bars, 20 μm. Graphs in lower panels show quantification of the indicated markers, normalized to the area of the primary lobe (*p < 0.05, **p < 0.01). Error bars represent SD. *TepIV* >, n = 8; *TepIV* > *Su(H)*^RNAi 1^, n = 5. **(D-E) Overexpression of Notch (D) or Supressor of Hairless (E) in Core Progenitors has no effect on cell differentiation.** Core Progenitors are visualized in green (*TepIV* > GFP); Plasmatocytes in red (P1 staining); and Crystal Cells in cyan (ProPO staining). Upper panels: Control lymph glands; middle panels: Lymph glands with *TepIV*-Gal4 driven overexpression of full-length Notch (**D**) or Su(H) (**E**). Whole Z-projections of confocal images of primary lobes from wandering third instar larvae are shown. The dashed line marks the outline of the lymph gland lobes. Scale bar, 50 μm for Notch (**D**) and 60 μm for Su(H) (**E**). Lower panels: Quantification of the results (ns: non-significant). Error bars represent SD. *TepIV* >, n = 8; *TepIV* > Notch, n = 8; *TepIV* > Su(H), n = 8. **(F) Schematic representation of the effect of Notch pathway manipulation in Core Progenitors**. Cartoon of a wandering 3^rd^ instar wild-type lymph gland lobe (cross section, anterior is up and center is to the left). Left: expression domain of *TepIV*, were the manipulations were performed (gray). Panels 2-4: core progenitors are colored in green, the Cortical Zone in pink, and Crystal Cells in cyan. Downregulation of Notch signaling in *TepIV* expressing cells provokes significant loss of Core Progenitors and increase of differentiated cells at the Cortical Zone. Notch signaling upregulation in Core Progenitors has no effect.

We next analyzed whether overactivation of the Notch pathway provokes the opposite phenotype, namely an increase of Core Progenitors and general differentiation impairment. This was not the case, as over-expression with *TepIV*-Gal4 of Notch (**Fig. S2A**) or Su(H) did not alter Core Progenitor, PL or CC populations (**Fig. 2D, E, F**). These results suggest that endogenous activity of the Notch pathway is already sufficient to prevent excessive differentiation of Core Progenitors. It was recently reported that Notch is expressed transiently at the 1^st^ larval instar (L1) in a small group of HSCs (Dey et al., 2016), so we analyzed the possibility that the phenotype that we observed in Core Progenitors stems from an alteration of Notch function in L1 HSCs. To investigate this, we utilized a thermosensitive Gal80 construct to silence Notch expression only from mid-second larval instar onwards (**Fig. S2F**, upper). The results were identical to those in which Notch was constitutively silenced (**Fig. 2B** and **Fig. S2B**), ruling out the possibility that the loss of Core Progenitors observed after Notch silencing depends on an early role in L1 HSCs. The findings shown in this section (**Fig. 2F**) are consistent with a requirement of the Notch pathway for maintenance of Core Progenitors in an undifferentiated state.

### Serrate expressed at the Posterior Signaling Center is required for Core Progenitor maintenance

As a next step, we studied the identity and source of the ligand for Notch activation in Core Progenitors. *Serrate* (*Ser*) is expressed at high levels in PSC cells (Ferguson and Martinez-Agosto, 2014; Jung et al., 2005; Lebestky et al., 2003; Mandal et al., 2007), although no functions have been yet attributed to this expression. Since the PSC and Core Progenitors are in close proximity, and filopodia that emanate from PSC cells may play a role in transmitting signals to MZ progenitors (Mandal et al., 2007), we asked whether *Ser* expressed at the PSC activates Notch for Core Progenitor maintenance. First, we silenced *Ser* at the PSC with an *Antp-*Gal4 driver, and observed an increased number of PLs and CCs (**Fig. 3A, Fig. S3A**), suggesting that Ser is required at the PSC to limit differentiation and to maintain Core Progenitors.

**Figure 3.**
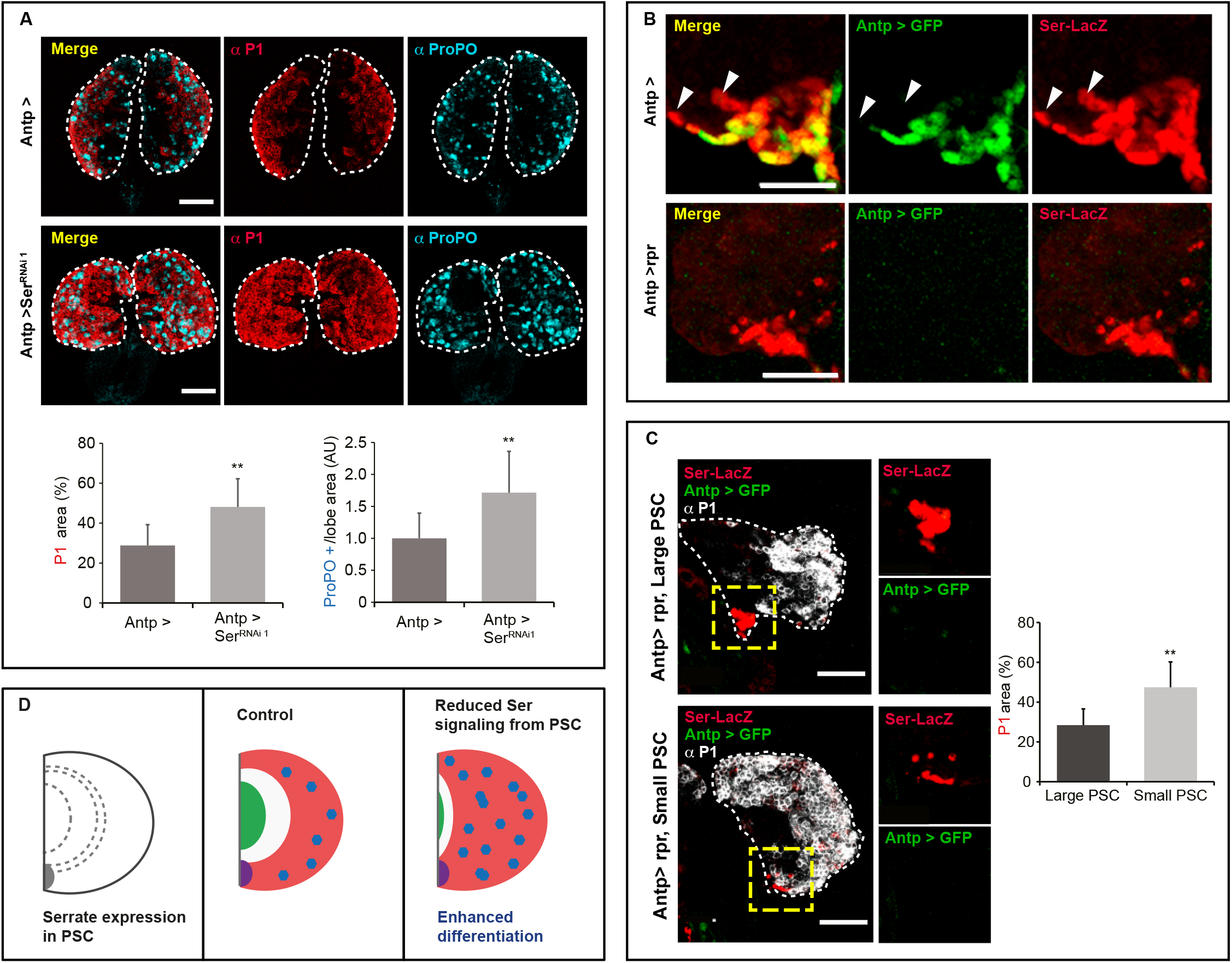
Serrate is required at the Posterior Signaling Center for Core Progenitor maintenance. **(A) *Serrate (Ser)* silencing at the Posterior Signaling Center provokes increased differentiation of Plasmatocytes and Crystal Cells.** Following *Serrate* RNAi (*Ser*^RNAi 1^) expression with an *Antp*-Gal4 driver, general increase of cell differentiation is observed. Plasmatocytes are shown in red (P1 staining) and Crystal Cells in cyan (ProPO staining). Upper panels: Control lymph glands; middle panels: Lymph glands where expression of *Ser*^RNAi 1^ was driven by *Antp*- Gal4. Whole Z-projections of confocal images of primary lobes from wandering third instar larvae are presented. The dashed line marks the outline of the primary lobes. Scale bars, 60 μm. Lower panels: Quantification of the results (**p < 0.01). Error bars represent SD. *Antp* >, n = 10; *Antp* > *Ser*^RNAi 1^, n = 14. **(B) A fraction of***Serrate***-expressing cells at the Posterior Signaling Center survive***Antennapedia***-driven genetic ablation.** *Antp*-Gal4 driven expression of GFP (*Antp* > GFP, green) at the PSC; *Serrate* (*Ser*) expressing cells are visualized by anti-βGal staining (red) of a *Ser*LacZ reporter. Expression of the pro-apoptotic protein Reaper (Rpr) with an *Antp*-al4 driver from L2 stage onwards (see text) eliminates all *Antp* > GFP cells, but many *Ser* expressing cells survive. Upper panels: control lymph glands without Rpr expression. The wild-type PSC comprises a mixed population of *Antp* > GFP plus *Ser* double-positive cells, and *Ser* positive plus *Antp* > GFP negative cells (examples of the latter are shown with arrowheads); lower panels: lymph glands expressing Rpr under an *Antp-*Gal4 driver. After genetic ablation, all the surviving PSC cells express *Ser* and lack *Antp > GFP* expression. Single-plane confocal images of the PSC region of primary lobes of wandering 3^rd^ instar larvae are shown. Scale bars, 30 μm. **(C) The number of***Serrate***-xpressing cells that survive genetic ablation correlates negatively with the amount of Plasmatocytes.** In lymph glands in which *Antp*-expressing cells of the PSC have been ablated by *Antp*-Gal4 driven expression of Rpr, Plasmatocyte differentiation (white, P1 staining) correlates negatively with the area occupied by cells expressing *Serrate*-LacZ that survived genetic ablation (red, βGal staining). Examples of large and small *Ser*positive PSC areas are shown (“Large PSC” on the upper panel and “Small PSC” on the lower panel). The inset shows a magnification of the PSC, with all PSC cells expressing *Ser* but not *Antp* > GFP. Whole Z-projection confocal images of primary lobes from wandering third instar larvae are depicted. The dashed line indicates the outline of the primary lobe. Scale bars, 20 μm. The graph shows quantification of P1-positive area in *Antp*-ablated lymph glands that exhibit large or small PSC area. PSC area was considered small if it was less than 1.5% of the total lobe area). Large PSC, n = 7, Small PSC, n = 8. **(D) Schematic representation of the results.** Cross sections of lymph gland lobes are represented. The PSC is highlighted in purple, Core Progenitors in green, the Cortical Zone in pink, and Crystal Cells in cyan. Reduced *Ser* signaling from the PSC (gray area in the left panel) due to *Ser* silencing or Rpr dependent ablation of PSC cells provokes an increase of differentiated cells at the Cortical Zone (compare center and right panels).

Recently, the notion that the PSC functions as a hematopoietic niche has been challenged, as *Antp*-Gal4 driven expression of reaper (i.e. apoptotic ablation of the PSC) did not alter PL or CC differentiation (Benmimoun et al., 2015). In that study, genetic ablation of the PSC was confirmed by the lack of expression of two classical PSC markers, *hh* and Antp, whereas *Ser* expression was not assessed (Benmimoun et al., 2015). We thus analyzed if *Ser* expressing cells are still present in lymph glands in which the PSC was genetically ablated. Ablation was induced from L2 stage onwards by using a Gal80 thermosensitive construct to prevent *reaper* expression and lethality at earlier stages (**Fig. S3B)**. We monitored the expression of a *Ser-LacZ* construct (Bachmann and Knust, 1998; Ferguson and Martinez-Agosto, 2014; Jung et al., 2005; Lebestky et al., 2003; Mandal et al., 2007), and consistently observed the presence of *Ser*-expressing cells in lymph glands, where cells positive for the *Antp*-Gal4/UAS-GFP marker were absent following genetic ablation (**Fig 3B,** lower). Importantly, in wild-type lymph glands we also observed PSC cells that express *Ser* but not the *Antp*-Gal4/UAS-GFP marker (**Fig. 3B,** upper), suggesting that the PSC might encompass a mixed population of cells, with a large proportion of them coexpressing *Antp* > GFP and *Ser*, while the remaining PSC cells express *Ser* but not *Antp* > GFP. These observations are consistent with a model in which, after *Antp*-Gal4 driven ablation of the PSC, the remaining cells of the PSC that express *Ser* but not *Antp*-Gal4 are sufficient to sustain normal Core Progenitor maintenance and hemocyte differentiation. Noteworthy, after *Antp* driven ablation of PSC cells, a negative correlation between surviving *Ser*^+^, *Antp*Gal4^-^ PSC cells and hemocyte differentiation occurs: Lymph glands with a small *Ser*-positive area at the PSC display greater PLs differentiation than those with a larger *Ser*-positive area at the PSC (**Fig. 3C**). These results suggest that *Ser* expressing cells of the PSC are important to restrain Core Progenitor differentiation (**Fig. 3D**), and that the *Ser* expressing cells that survive to *Antp*-driven apoptotic cell ablation can support normal functions of the PSC.

We analyzed in more detail in wild-type lymph glands those PSC cells that express *Ser* but are negative for *Antp*-Gal4, and found that they all stain positive with anti-Antp (**Fig. 4A)** and anti- Collier antibodies (**Fig. S3C)**, while they all express the *hh*-GFP marker as well (**Fig. S3C)**. These observations suggest that this population of *Ser*-expressing cells belongs entirely to the PSC, which was confirmed by their lack of expression of *domeMESO* > GFP **(Fig. 4B)**. The above results imply that the *Antp* enhancer that controls Gal4 expression is not active at late 3^rd^ larval instar in a subpopulation of PSC cells. To better characterize this cell heterogeneity within the PSC, we performed a cell lineage analysis by utilizing the G-TRACE system driven by *Antp*-Gal4. We found that all the PSC cells express the lineage tracer GFP, while a subpopulation fails to express the real time expression marker RFP under *Antp*-Gal4 at late 3^rd^ larval instar (**Fig. 4C)**. These results indicate that all PSC cells derive from an original *Antp*-Gal4 positive lineage, although some of them have turned off its expression. Finally, we confirmed that *Antp*-Gal4 driven expression of Reaper mediates ablation of just a subpopulation of PSC cells, while the surviving cells that express *Ser* are immunoreactive for Antp (**Fig. 4D, lower)**, and therefore belong to the PSC. In contrast, expression of *reaper* under control of a *col*-Gal4 driver led to total ablation of the PSC, as revealed by the absence of cells that stain positive with anti-Antp antibody, as well as the lack of *hh*-GFP or *Ser*-LacZ expressing cells (**Fig. 4E; Fig. S3D; Fig. S3E)**. These lymph glands in which the PSC has been completely ablated display enhanced PL and CC differentiation (**Fig. 4F),** supporting again the notion that the PSC is important to restrain differentiation and for maintenance of progenitors in an undifferentiated state (**Fig. 4G)**(Mandal et al., 2007; Mondal et al., 2011; Pennetier et al., 2012).

**Figure 4.**
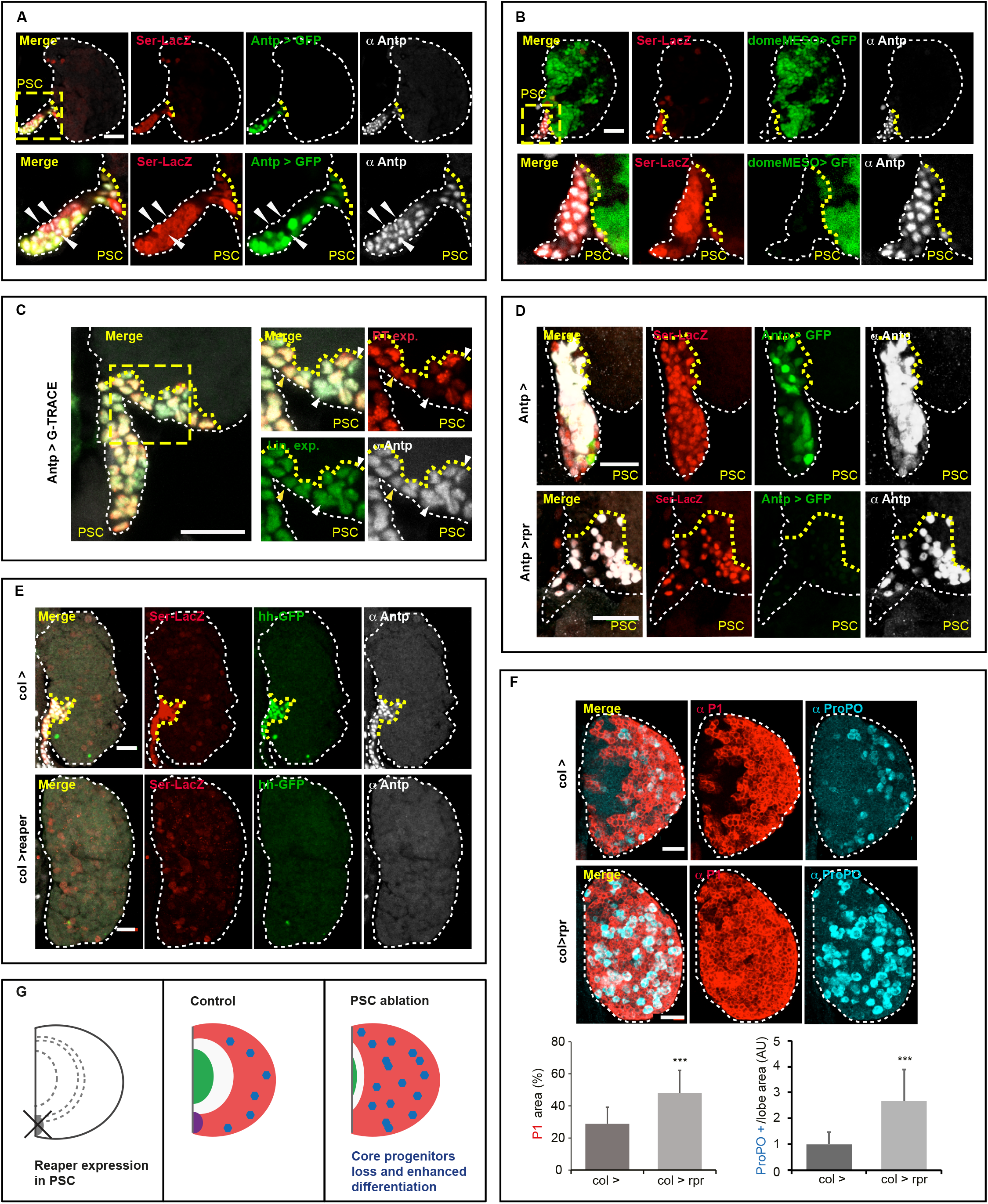
The Posterior Signaling Center is required for Progenitor maintenance and to limit cell differentiation. **(A) The PSC is composed of a heterogeneous population of cells.** While all cells of the PSC (marked with a yellow square and zoomed in the lower row) express *Ser*-LacZ and stain positive with an anti-Antp antibody, some cells do not express GFP driven by *Antp*-Gal4 (examples marked with arrowheads). The white dashed line indicates the outline of the primary lobe. Yellow lines mark the limit of the PSC. Single-plane confocal images of the PSC region of primary lobes from wandering 3^rd^ instar larvae are shown. Scale bars, 20 μm. **(B) The *domeMESO* GFP reporter is not expressed in the Posterior Signaling Center cells.** The square indicates the position of the PSC, magnified in lower panels. PSC cells are visualized with an anti-Antp antibody (white), and by *Ser*-LacZ expression (red), while progenitors of the Medullary Zone express the *domeMESO* > GFP reporter (green). Note that *Ser*-positive cells do not coexpress *domeMESO*. The white dashed line indicates the contour of the primary lobe. The yellow line marks the PSC border. Single-plane confocal images of the PSC from wandering third instar larvae are shown. Scale bars, 20 μm. **(C) All PSC cells derive from an***Antp***-Gal4 positive lineage.** G-TRACE analysis performed with an *Antp*-Gal4 driver. The GFP signal marks the *Antp* cell lineage (green), while the RFP signal indicates ongoing expression of *Antp* at the moment of the experiment (red). PSC cells are stained with an anti-Antp antibody (white). The white dashed line marks the contour of the primary lobe, and the yellow line, the limit of the PSC. The zoomed in region is indicated with a yellow line on the left panel. Whole Z-projections of confocal images of the PSC from wandering third instar larvae are shown. Scale bars, 20 μm. **(D) The cells that survive***Antp***-Gal4 driven ablation belong to the PSC.** Whole Z-projection of confocal planes of the PSC, where genetic ablation was induced by expressing *UAS-rpr* with an *Antp*-Gal4 driver (lower). The surviving cells express *Ser*-Lac (red) and are decorated with Antennapedia protein (white), but fail to express the *Antp*-Gal4 > GFP reporter (green). The dashed line marks the outline of the primary lobe. The yellow line indicates the PSC border. Scale bars, 20 μm. **(E) The PSC was fully ablated after***collier***Gal4 driven expression of***reaper*. Upper panels (*col*>): Control lymph glands with a normal PSC; lower panels: Lymph glands expressing *rpr* under control of a *col*-Gal4 driver. In control individuals, lymph glands display expression at the PSC of the reporters *Ser*-Lac (red) and *hedgehog*-GFP (green), and stain positive with an α-Antp antibody (white). Upon *col*-Gal4 dependent genetic ablation, all PSC cell were eliminated, so none of the PSC markers can be detected. Whole Z-projections of confocal images of the PSC region of primary lobes from wandering 3^rd^ instar larvae are shown. The white dashed line marks the outline of the primary lobe. The yellow line marks the PSC limit. Scale bars, 20 μm. **(F) *collier* Gal4 driven genetic ablation of the PSC induces Plasmatocyte and Crystal Cell differentiation.** Whole Z-projections of primary lobes where Plasmatocytes (red: P1 staining) and Crystal Cells (cyan: ProPO staining) are visualized. Upper panels: Control lymph glands; middle panels: Lymph glands with *collier-Gal4* induced expression of *rpr*. The dashed line marks the outline of the primary lobe. 20 μm. Lower panels: Charts show quantification of the indicated markers (***p < 0.001). Error bars represent SD. *col* >, n = 18; *col* > *rpr*, n = 19. **(G) Schematic representation of the results.** Cross sections of lymph gland lobes are represented (anterior is up and center is left). The PSC is highlighted in purple, the Core Progenitors in green, the Cortical Zone in pink, and the Crystal Cells in cyan. *collier*-Gal4 driven expression of *reaper* in the PSC (gray area in the left panel) reduces the amount of progenitors and induces Plasmatocyte and Crystal Cell differentiation (right panel).

### In Distal Progenitors the Notch pathway controls a binary cell fate decision

We have shown above that *Notch* silencing with *Tep-IV*Gal4 provokes enhanced differentiation of both PLs and CCs (**Fig. 2B**). In sharp contrast, *Notch* silencing with *domeMESO*-Gal4 brought about almost complete loss of CCs accompanied by an increase of PLs (**Fig. 5A; Fig. S4A**). Similar results were obtained when *Su(H)* was silenced with the same driver (**Fig. 5B; Fig. S4B**), suggesting that the Notch pathway promotes CC differentiation in Distal Progenitors, while the PL fate is inhibited. To further explore this possibility, we over-expressed with *domeMESO*-Gal4 a full-length Notch construct (**Fig. S4C**) or the Notch Intracellular Domain (NICD), and observed in both cases that PL differentiation was virtually blocked, and CCs increased dramatically (**Fig. 5C**). Overexpression of Su(H) with *domeMESO*-Gal4 provoked the same effect (**Fig. 5D**). We thus conclude that an increase of Notch pathway activity in Distal Progenitors induces CC and inhibits PL differentiation **(Fig 5E)**.

**Figure 5.**
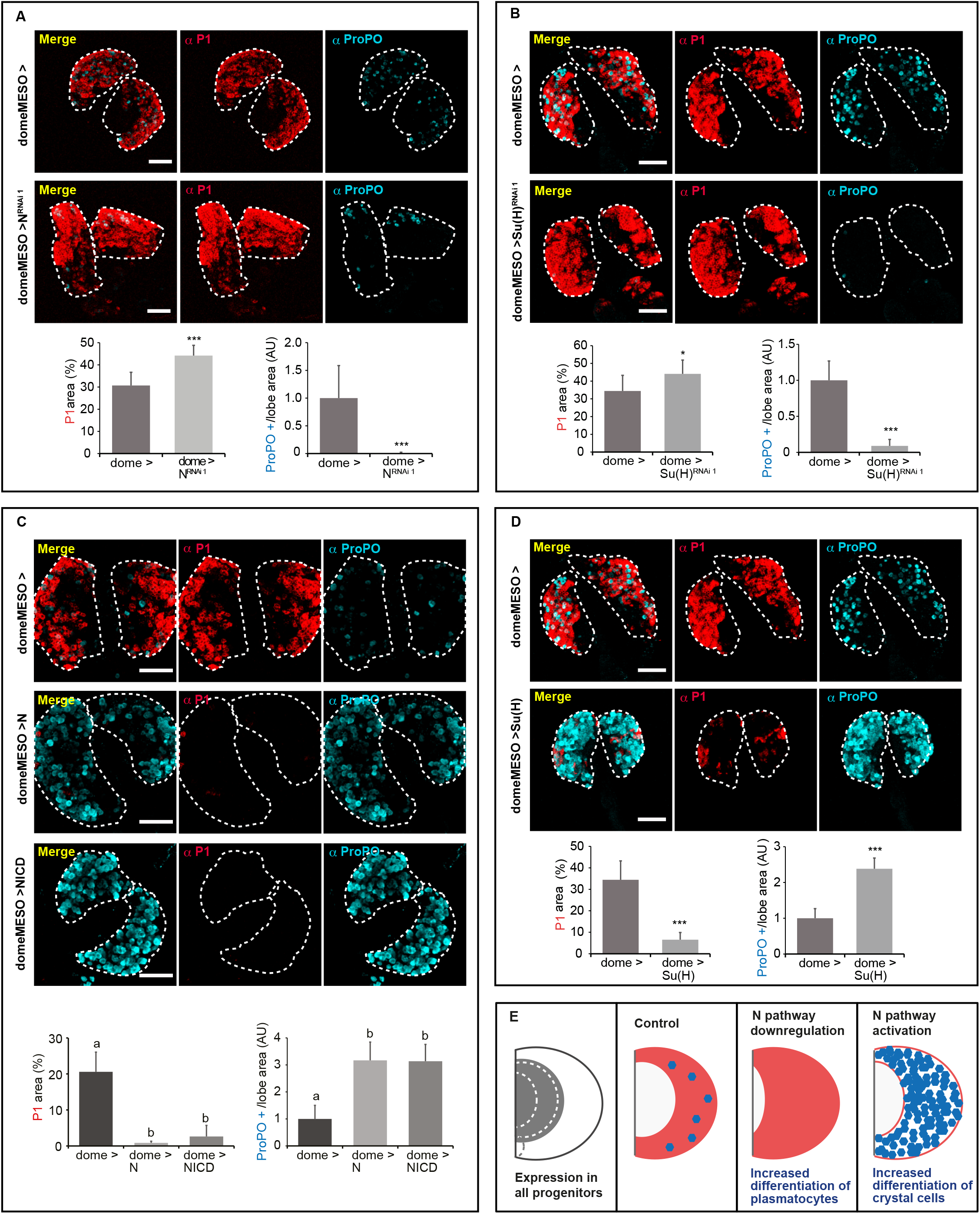
Notch regulates the Plasmatocyte/Crystal Cell fate decision in Distal Progenitors. **(A) *Notch* silencing in *domeMESO*-positive progenitors abolishes Crystal Cell specification and promotes the Plasmatocyte cell fate.** Whole Z-projections of confocal images of primary lobes of wandering 3^rd^ instar larvae stained for P1 (red, Plasmatocytes) and ProPO (cyan, Crystal Cells) are displayed. Compare control lymph glands in upper panels with lymph glands expressing *N^RNAi1^* in middle panels. The dashed lines mark the outline of primary lobes. Scale bars, 60 μm. Charts in lower panels show quantification of the indicated markers (***p < 0.001). Error bars represent SD. *domeMESO* >, n = 8; *domeMESO* > N^RNAi 1^, n = 8. **(B) *Suppressor of Hairless* silencing in *domeMESO*-positive progenitors provokes a similar effect to that of *Notch* silencing.** *Su(H)* RNAi (*Su(H)^RNAi 1^*) expression driven by *domeMESO*-Gal4 provoked loss of Crystal Cells (cyan: ProPO staining) and increased Plasmatocytes (red: P1 staining) (Middle panels). The dashed line marks the outline of the two lymph gland lobes. Scale bars, 40 μm. Whole Z-projections of confocal images of primary lobes of wandering 3^rd^ instar larvae are shown. Lower panels: quantification of the indicated markers (*p < 0.05, ***p < 0.001). Error bars represent SD. *domeMESO* >, n = 10; *domeMESO* > *Su(H)^RNAi 1^*, n = 14. **(C) Increased Notch activity in***domeMESO***-positive progenitors favors the Crystal Cell fate while inhibiting the Plasmatocyte fate.** In upper panels, a control lymph gland is shown. Below, lymph glands overexpressing full-length Notch (2^nd^ row) or NICD (3^rd^ row) under control of a *domeMESO*-Gal4 driver are shown. The dashed line indicates the outline of the primary lobes. Whole Z-projections of confocal images of primary lobes of wandering 3^rd^ instar larvae are depicted. Scale bars, 60 μm. Bottom row: Quantification of the results. Different letters indicate statistical difference (Tukey′s multiple comparison test). Error bars represent SD. *domeMESO* >, n = 6; *domeMESO* > Notch, n = 6; *domeMESO* > NICD, n = 4. **(D) Overexpression of Suppressor of Hairless mimics overexpression of Notch.** Whole Z- projections of confocal images of primary lobes of wandering 3^rd^ instar larvae are displayed. Plasmatocytes are visualized in red (P1 staining), and Crystal Cells in cyan (ProPO staining). Upper panels: Control lymph glands; middle panels: Lymph glands with *domeMESO*-Gal4 driven overexpression of Su(H). The dashed lines stand out the contour of primary lobes. Scale bars, 60 μm. Lower panels: Quantification of the results (***p < 0.001). Error bars represent SD. *domeMESO* >, n = 7; *domeMESO* > Su(H), n = 8. **(E) Summary of the results of Notch Pathway manipulations in *domeMESO* positive progenitors.** Schematic representation of a lymph gland primary lobe (cross section, anterior is up and center is left). Plasmatocytes are represented in pink and Crystal Cells in cyan. Left: *domeMESO* driver expression area (gray). *Notch* or *Su(H)* silencing in the *domeMESO* domain completely blocks Crystal Cell differentiation and provokes increase of Plasmatocytes localized at the Cortical Zone. Conversely, *domeMESO*-Gal4 driven Notch or Su(H) overexpression increases Crystal Cell differentiation and virtually blocks the Plasmatocytes fate.

To confirm that the activity of Notch is indeed required for normal differentiation in Distal Progenitors that are positive for *domeMESO* and negative for *Hml* (**Fig. 1D,** region 2), *Notch* silencing with *domeMESO*-Gal4 was repeated in a genetic background in which *Notch* RNAi expression in cells that coexpress *Hml* was inhibited by *QUAS-*Gal80 (*domeMESO-*Gal4 *> UAS N^RNAi^; Hml*-QF > *QUAS-* Gal80). Silencing of *Notch* in this genetic background provoked identical effects to those observed without expression of Gal80 in *Hml*-positive cells (**Fig. S4D**). In a control experiment, *Hml*-QF driven expression of *QUAS*-Gal80 effectively repressed Gal4 activity in the CZ (**Fig. S4E**). These results confirm that Notch has a function in Distal Progenitors, which express *domeMESO* but not *Hml*. Taken together, these experiments indicate that the Notch pathway regulates a binary fate decision in Distal Progenitors, promoting CC differentiation, while inhibiting the PL fate **(Fig 5E)**.

### eater is an early Plasmatocyte fate marker in Distal Progenitors

Next, we explored further aspects of this Notch-dependent binary fate decision that takes place in Distal Progenitors. The gene *eater* is expressed in PL and also in lymph gland progenitors (Kocks et al., 2005; Kroeger et al., 2012). Careful analysis of the expression of an *eater*-dsRed reporter revealed that it can be detected in most Distal Progenitors, but is virtually absent in a few of them (**Fig. 6A**). A possible explanation is that *eater*-expressing Distal Progenitors at the 3^rd^ larval instar might be already committed towards a PL fate, while those Distal Progenitors with very low *eater* levels are likely committed to become CCs. Consistent with this possibility, we observed at the CZ that *eater*-dsRed expression occurs in most *Hml*-positive cells, while it is excluded from those cells that express the CC progenitor marker lozenge (Lz) (**Fig. 6B**). Lineage tracing analysis with an *eater*-Gal4 driver confirmed that the *eater* lineage does not include CCs (**Fig. 6C**), supporting that *eater* expression in progenitors marks a Distal Progenitor subpopulation committed for a PL cell fate. *eater*-Gal4 driven silencing or over-expression of *Notch* did not recapitulate the effects observed when *domeMESO*-Gal4 was utilized to perform the same manipulations (compare **Figs. 5A, C** with **Fig. S5**), suggesting that Notch operates in Distal Progenitors that have not yet begun to express *eater*. The above observations are consistent with a model in which, at an earlier developmental stage, *eater* negative uncommitted Distal Progenitors make the binary cell fate decision regulated by Notch, while later, *eater* is expressed in cells committed to a PL fate. If *eater* is indeed an early marker for Distal Progenitors committed to a PL fate, its expression might be controlled by the Notch pathway. We analyzed this possibility by overexpressing Notch with *domeMESO-* Gal4, a treatment that induces massive differentiation of CCs (**Fig. 5C**), and observed almost complete absence of *dome*^+^, *eater*^*+*^ double-positive Distal Progenitors (**Fig. 6D**). These observations indicate that *eater* expression is repressed by the Notch pathway in uncommitted Distal Progenitors.

**Figure 6.**
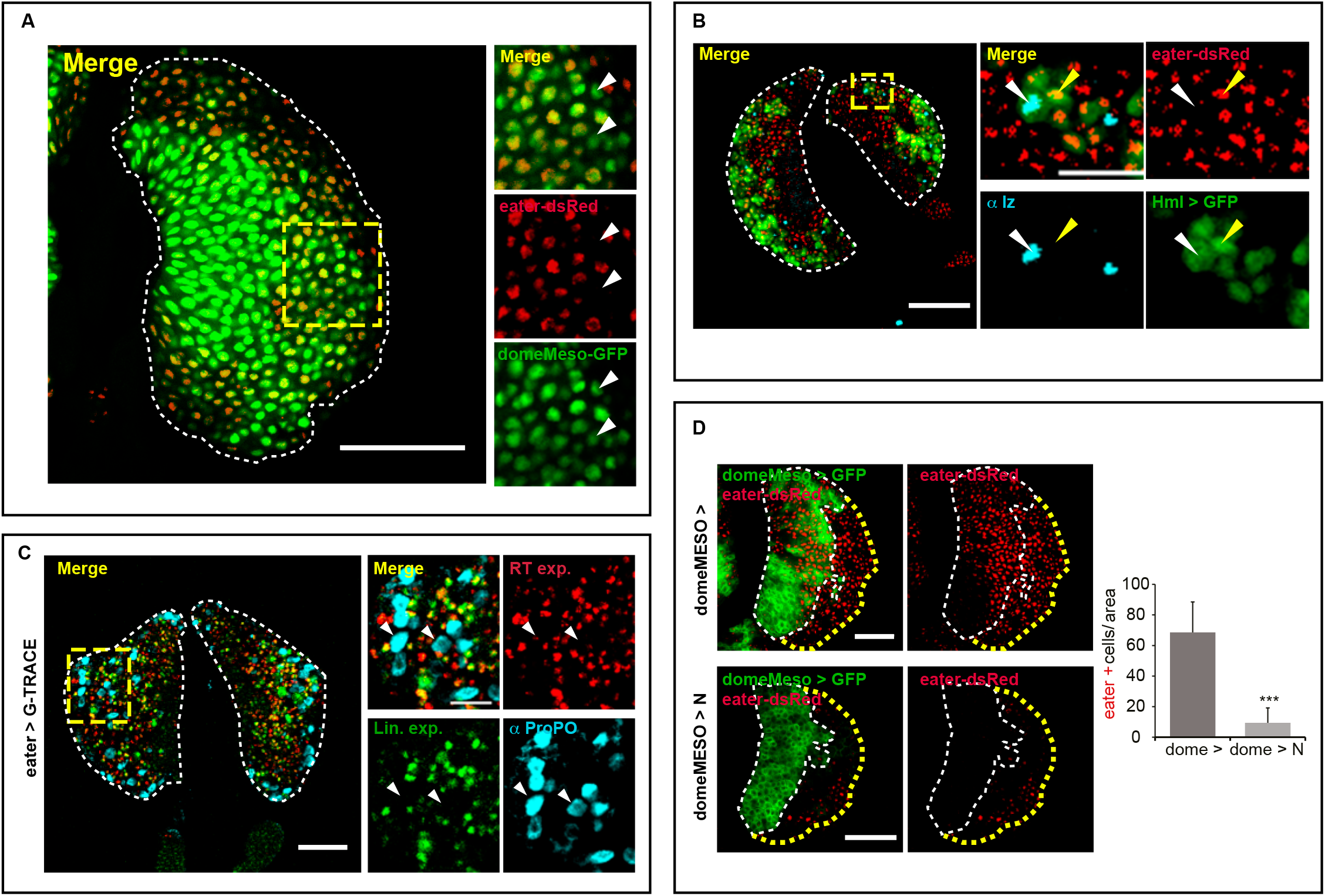
*eater* is an early Plasmatocyte fate marker in Distal Progenitors. **(A) *eater*-dsRed is expressed in most but not all lymph gland progenitors.** Left: Single-plane confocal image of a primary lobe of a 3^rd^ instar larva lymph gland, showing expression of the reporters *domeMESO*-GFP (green) and *eater*-dsRed (red). Right: Magnified image of the area marked with a yellow square, revealing that *eater*-dsRed is expressed at high levels in most Distal Progenitors, while it is barely detectable in a few cells (white arrowheads). The dashed line marks the outline of the lymph gland primary lobe. Scale bar, 20 μm. **(B) Crystal Cell Precursors do not express***eater*. Anti-Lozenge (Lz) staining (cyan, Crystal Cells) and *eater*-dsRed expression (red) are shown in lymph glands that also express GFP under *Hml*- Gal4 (green). In the magnified area, marked with the yellow rectangle, an example of an *Hml* > GFP positive cell that expresses the CC precursor marker Lz, but not *eater* (white arrowhead) is shown. The yellow arrowhead marks an example of the converse situation in which an *Hml* > GFP positive cell expresses *eater*-dsRed and does not stain positive for Lz. The dashed lines indicate the contour of primary lobes. Single-plane confocal images of primary lobes from wandering 3^rd^ instar larvae are shown. Scale bar, 40 μm. **(C) *eater* is not expressed during Crystal Cell differentiation.** An *eater*-Gal4 driver was utilized to express the G-TRACE system and the cell lineage is labeled with GFP (green), while real-time expression of *eater*-Gal4 is labeled with RFP (red). Crystal Cells are visualized by anti-ProPO staining (cyan), and display neither real time (RT exp., red) nor lineage expression (Lin exp., green) of the *eater*-Gal4 driver (two examples marked by arrowheads). The dashed line marks the outline of the two lymph gland lobes. Single-plane confocal images of primary lobes of wandering 3^rd^ instar larvae are shown. Scale bar, 50 μm. **(D) Notch represses *eater* expression in *domeMESO*-positive cells.** *eater*-dsRed reporter expression (red) is shown in control lymph gland (upper panels), and in lymph glands where over-expression of Notch was induced by *domeMESO*-Gal4 (lower panels). *domeMESO*-positive cells are revealed by expression of GFP (green). The yellow dashed line indicates the outline of the primary lobe. The white dashed line marks the *domeMESO* > GFP territory. Single-plane confocal images of primary lobes from wandering 3^rd^ instar larvae are shown. Scale bar, 40 μm. The chart shows quantification of double positive *domeMESO*^+^, *eater*^+^ cells in control and Notch overexpressing lymph glands (***p < 0.001). Error bars represent SD. *domeMESO* >, n = 9; *domeMESO* > N, n = 9.

### Serrate expressed in Lineage Specifying Cells at the MZ/CZ boundary regulates the cell fate decision of Distal Progenitors

Lineage Specifying Cells (LSCs) are a group of *Ser*-expressing cells that do not belong to the PSC, localize next to the MZ/CZ boundary, and instruct progenitors to acquire a CC fate (Ferguson and Martinez-Agosto, 2014; Lebestky et al., 2003). We therefore investigated whether these cells provide the ligand for the Notch-dependent binary cell fate decision in Distal Progenitors. As expected, these *Ser*-expressing cells, co-express *domeMESO* but not the PSC marker Antp **(Fig. 7A)**, so we utilized *domeMESO*-Gal4 to manipulate *Ser* expression. Silencing of *Ser* with *domeMESO*-Gal4 virtually blocked CC differentiation, while it increased the PL proportion (**Fig. 7B; Fig. S6A**), mimicking the effect of *Notch* silencing with the same Gal4 driver (**Fig. 5A**). In line with this, *domeMESO*-Gal4 driven overexpression of Ser rendered the opposite effect, increasing CCs and reducing PLs (**Fig. 7C**). Thus Ser expressed in LSCs is required for Notch activation in Distal Progenitors, controlling the binary cell fate choice. The E3 ubiquitin ligase Neuralized (Neur) is necessary for Ser endocytosis and Notch activation in neighboring cells (Le Borgne et al., 2005). Mimicking the results obtained after *Ser* silencing with *domeMESO-*Gal4, *neur* RNAi-mediated silencing with the same driver reduced the CC number and increased PLs (**Fig. 7D; Fig. S6B**), further supporting the notion that Ser expressed in LSCs activate Notch signaling for the binary cell fate decision of the Distal Progenitors **(Fig. 7E)**.

**Figure 7.**
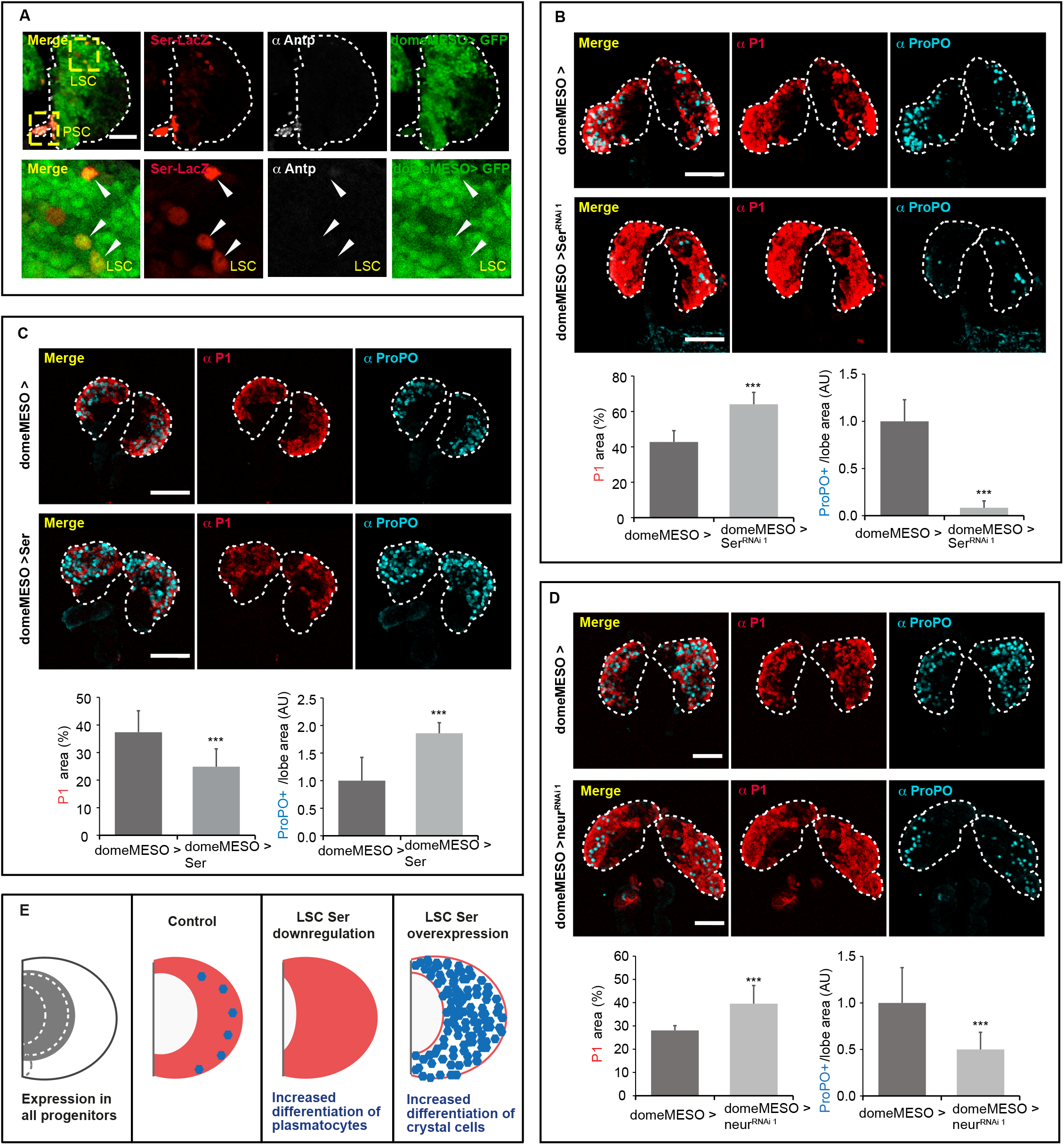
*Serrate* expression in Lineage Specifying Cells regulates the Plasmatocyte/Crystal Cell fate decision in Distal Progenitors. **(A) Lineage Specifying Cells coexpress *Ser* and *domeMESO*.** *Ser*-positive cells (red) of the PSC (yellow square) express Antp (white) and do not express the *domeMESO* > GFP reporter (green; upper panels). *Ser*-positive LSCs (yellow square in left upper panel) do not express Antp and coexpress *domeMESO.* Lower panels depict magnified images of the LSCs marked with the yellow square. Arrowheads mark examples of LSCs, which are negative for Antp and positive for domeMESO. Single-plane confocal images of primary lymph gland lobes of mid-3^rd^ instar larvae are shown. Scale bar, 40 μm. **(B) *Serrate* at the Medullary Zone is required for Crystal Cell differentiation and Plasmatocyte fate inhibition.** Plasmatocytes are visualized in red (P1 staining) and Crystal Cells in cyan (ProPO staining) in control lymph glands (upper panels) and in lymph glands that express a *Ser RNAi* (*Ser^RNAi 1^*) under *domeMESO*-Gal4 driver control (middle panels). Note that *Ser* silencing in LSCs leads to reduced CCs and increased PLs. The dashed line indicates the outline of primary lobes. Whole Z-projections of confocal images of primary lobes from wandering third instar larvae. Scale bar, 40 μm. Lower panels: Quantification of the results (***p < 0.001). Error bars represent SD. *domeMESO* >, n = 10; *domeMESO* > *Ser*^RNAi 1^, n = 12. **(C) Over-expression of Serrate (Ser) at the Medullary Zone provokes increased differentiation of Crystal Cells and reduces Plasmatocytes.** Plasmatocytes (red, P1 staining) and Crystal Cells (cyan, ProPO staining) in control lymph glands (upper panels) and in lymph glands with *domeMESO*-Gal4 driven Ser overexpression (middle panels). Note that Ser overexpression provokes an increase of CCs and a decrease of PLs. The dashed line marks the limits of the lymph gland primary lobes. Whole Z-projections of confocal images of primary lobes from wandering 3^rd^ instar larvae are shown. Scale bar, 50 μm. Lower panels: Quantification of the results (***p < 0.001). Error bars represent SD. *domeMESO* >, n = 12; *domeMESO* > Ser, n = 12. **(D) Neuralized (Neur) at the Medullary Zone is required for Crystal Cell differentiation and Plasmatocyte fate inhibition.** Plasmatocytes (red, P1 staining) and Crystal Cells (cyan, ProPO staining) are shown in control lymph glands (upper panels) and in lymph glands that express *neur* RNAi (*neur*^RNAi1^) under *domeMESO*-Gal4 (middle panels). Note that *neur* silencing provoke PL increase and CC reduction. The dashed line marks the outline of lymph gland lobes. Whole Z-projections of confocal images of primary lobes of wandering 3^rd^ instar larvae are shown. Scale bar, 40 μm. Lower panels: Quantification of the indicated markers (***p < 0.001). Error bars represent SD. *domeMESO* >, n = 10; *domeMESO* > *neur*^RNAi 1^, n = 10. **(E) Schematic representation of the results.** Cartoon of a primary lobe of a 3^rd^ instar lymph gland primary lobe. *Serrate* manipulation at the *domeMESO* territory (gray region in left panel) regulates the fate that Distal Progenitors will adopt. *Ser* silencing with *domeMESO*-Gal4 provokes an increase of Plasmatocytes (pink) and blocks Crystal Cell differentiation (cyan). Serrate overexpression driven by *domeMESO*-Gal4 induces Crystal Cells blocks Plasmatocyte differentiation.

## Discussion

In mammals, Notch receptors (Notch 1-4) are expressed in HSCs, hematopoietic progenitors and mature blood cells, suggesting that Notch is required at multiple levels of blood cell differentiation. Particularly, the role of the Notch pathway in HSC maintenance has been explored in depth, although with contrasting results. Some experiments in cell culture suggest that Notch is required for HSC maintenance (Kumano et al., 2001; Varnum-Finney et al., 2000; Vercauteren and Sutherland, 2004), while others suggest that it promotes HSC differentiation towards the myeloid lineage (Schroeder and Just, 2000; Schroeder et al., 2003). Works *in vivo* in which interactions with the hematopoietic niche are still present, and therefore recapitulate better a physiologic situation, also yielded conflicting results. Studies utilizing conditional knock-out mice affecting different elements of the canonical Notch pathway suggested that Notch is dispensable for HSC maintenance (Maillard et al., 2008; Mancini et al., 2005), while others indicated that alterations of Notch signaling are associated with development of myeloproliferative disease (MPD), characterized by accumulation of mature cells of the myeloid lineage (Klinakis et al., 2011). Thus, the role of Notch in adult HSC homeostasis is controversial and requires further examination. Many aspects of mammalian and *Drosophila* hematopoiesis are conserved, particularly at the level of the transcription factors and signaling pathways involved (Banerjee et al., 2019; Crozatier and Vincent, 2011; Evans et al., 2007). One striking parallelism between *Drosophila* and mammalian models is the expression in the hematopoietic niche of the Notch ligand Serrate (Ser)/Jagged (JAG). While Ser is expressed in cells of the *Drosophila* PSC (Lebestky et al., 2003), mammalian JAG1 and JAG2 expression has been detected in different bone marrow cell types that fulfill niche functions, including endothelial cells and cells of the hematopoietic stroma (Calvi et al., 2003; Fernandez et al., 2008; Varnum-Finney et al., 1998). The wide array of genetic tools available in *Drosophila* allow for manipulations of gene expression with an exquisite temporal and spatial specificity that is currently not possible to the same extent in mammalian systems. Thus, the utilization of the fly model may provide clues for addressing unresolved issues related to Notch functions in mammalian hematopoiesis.

In the current work, we have analyzed biological properties of *Drosophila* blood cell progenitors. This analysis led us to redefine the progenitor subpopulations of the larval lymph gland. A novel progenitor population, the “Distal Progenitors” that are positive for *domeMESO* and negative for both *TepIV* and *Hml*, occurs at the MZ. Our results strongly suggest that the binary fate decision between PLs and CCs is made in naïve Distal Progenitors at early larval stages, while later in development, at wandering 3^rd^ instar larvae, the gene *eater* is expressed in most but not all Distal Progenitors, marking the cells that are already committed to a PL fate. The remaining Distal Progenitors with low *eater* levels are committed to become CCs, as suggested by *eater* lineage tracing, as well as by *eater* and Lz mutually exclusive expression in *Hml*-positive cells. Thus, the MZ, in wandering 3^rd^ instar larvae, encompasses an undifferentiated population of cells, the Core Progenitors, and a second population, the Distal Progenitors, which have already been instructed to become either PLs or CCs (**Fig. 8**).

**Figure 8.**
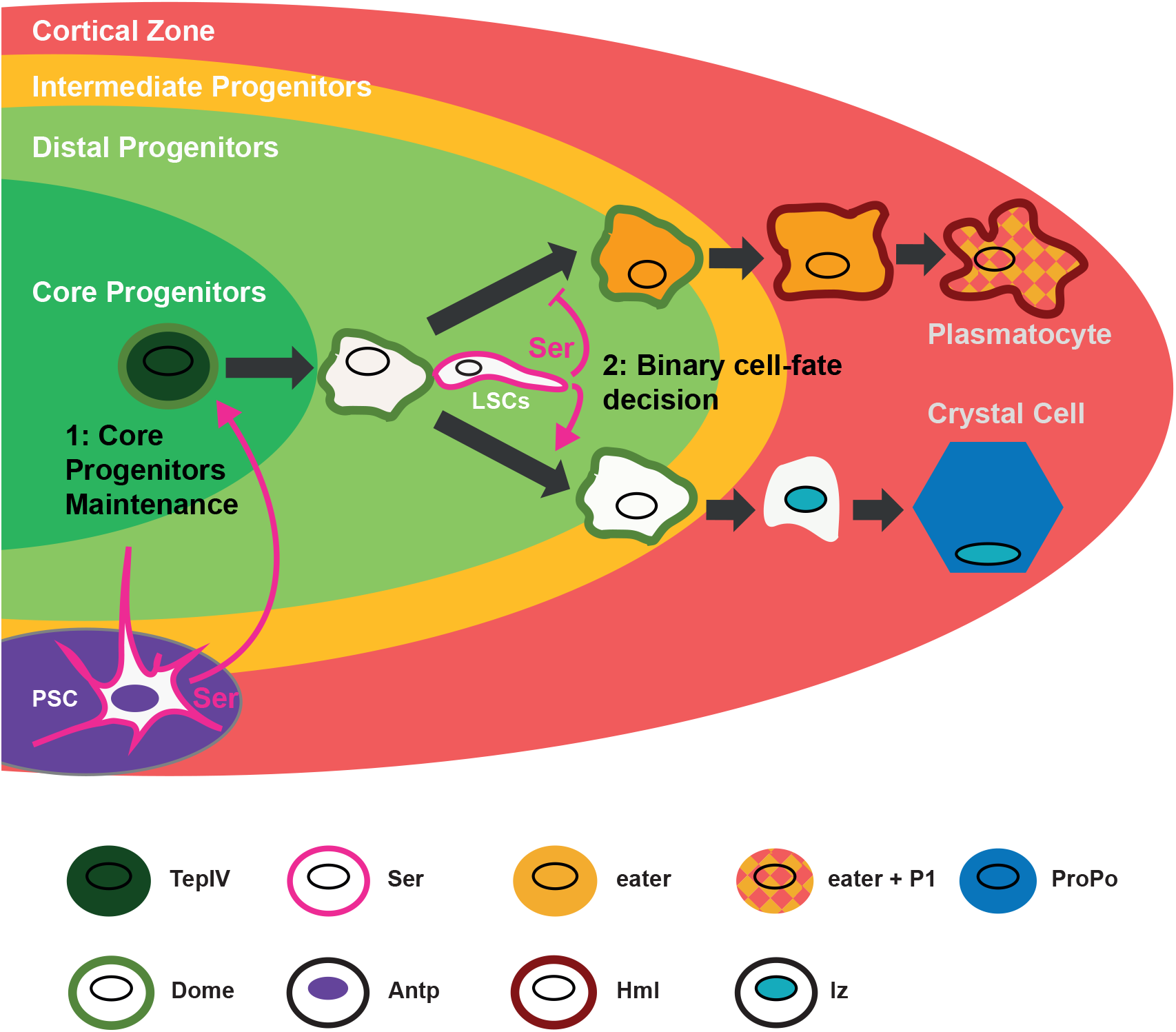
Model for the function of Notch in the regulation of lymph gland differentiation. In *TepIV^+^ domeMESO^+^* Core Progenitors (dark green) Notch is activated by its ligand Ser expressed at the PSC. This Notch activation is required to avoid premature Core Progenitor differentiation to Plasmatocytes and Crystal Cells (1: Core Progenitors maintenance). Thereafter *TepIV* expression ceases, while *domeMESO* expression continues, defining a second population of progenitors, the Distal Progenitors (light green). Early Distal Progenitors do not express any differentiation marker, and constitute the stage where a cell fate decision between the Plasmatocyte and the Crystal Cell fate is made. This cell fate decision also depends on Notch (2: Binary cell fate decision), activated by Ser expressed in the Lineage Specifying Cells (LSCs), which coexpress *domeMeso*. Notch activation in Distal Progenitors induces CC differentiation and inhibits the Plasmatocyte fate. Without contact with Ser-expressing LSCs, Distal Progenitors follow a default Plasmatocyte differentiation program. This early fate decision is evidenced later in development by the expression of *eater* in the Distal Progenitors committed to a Plasmatocyte fate (*eater* expression is represented with an orange background). Once at the Cortical Zone (pink), cells committed to a Plasmatocyte fate express hemolectin and continue expressing *eater*, while cells committed to become Crystal Cells now express Lozenge (Lz, light blue). Finally, mature PLs coexpress *eater*, *Hml* and the P1 antigen; while mature CCs coexpress Lz and ProPO (dark blue).

We have found that Notch, which is expressed throughout the lymph gland, plays distinct functions in Core Progenitors or Distal Progenitors (**Fig. 8**). In Core Progenitors Notch is required for maintenance of the undifferentiated state; our results thus contribute to the incipient characterization of the Core Progenitor subpopulation, in which Collier (Col) is expressed at low but physiologically relevant levels (Oyallon et al., 2016). We found that Notch function in Core Progenitors depends on Ser that is expressed at the PSC. This Ser expression has been reported before (Ferguson and Martinez-Agosto, 2014; Jung et al., 2005; Lebestky et al., 2003; Mandal et al., 2007), although its function has remained elusive. Filopodia that emanate from PSC cells and intermingle between cells of the MZ have been described, and were proposed to participate in Hedgehog signaling (Mandal et al., 2007). Given our observation that Ser expressed at the PSC is necessary for Notch activation in Core Progenitors, and considering that direct cell-cell interactions are required for Notch stimulation by its ligands, it seems reasonable to hypothesize that the filopodia that emanate from the PSC may play a role in Notch signaling as well (Krzemien et al., 2007).

It has been reported that genetic ablation of the PSC by expression of the proapoptotic protein Reaper (Rpr) does not alter Core Progenitor maintenance or steady-state differentiation, challenging the notion that the PSC functions as a hematopoietic niche (Benmimoun et al., 2015). Using a similar PSC-ablation protocol with *Antp*-Gal4 driven expression of Rpr from L2 stage onwards, we detected the presence of *Ser*-positive cells in the PSC that escaped genetic ablation. We found that this is because the *Antp*-Gal4 construct is not expressed in all PSC cells at late 3^rd^ larval instar, and thus that a subpopulation of PSC cells survive to Reaper-dependent ablation. The surviving PSC cells, which express *Ser* along with the other PSC markers, can support normal blood cell differentiation and core progenitor maintenance. When we utilized a *col*-Gal4 driver to express *reaper*, PSC cells were totally ablated and massive differentiation of Plasmatocytes and Crystal Cells was observed. Taken together, these observations might suggest that the PSC is necessary for restraining progenitor differentiation. However, given that, besides its strong expression at the PSC, *col-*Gal4 displays low expression levels in Core Progenitors (Crozatier et al., 2004; Oyallon et al., 2016), we cannot exclude the possibility that part of the effect of *col-*Gal4 dependent ablation on cell differentiation originates from its faint expression at the MZ.

We have found that in Distal Progenitors Notch fulfills a totally different function; it regulates a binary cell fate decision, promoting CC differentiation, and inhibiting differentiation of PLs (**Fig. 8**). We propose that the activation of the Notch pathway in Distal Progenitors is achieved through direct interaction of early naïve Distal Progenitors with *Ser*-expressing LSCs localized at the MZ/CZ boundary, thereby inducing CC differentiation and repressing the PL fate. On the other hand, naïve Distal Progenitors that do not contact these LSCs undergo a default differentiation program towards a PL fate (**Fig. 8**). Our finding that Notch regulates in Distal Progenitors a binary cell fate decision between a PL and a CC fate is in line with previous results by Tokusumi et al. (Tokusumi et al., 2010): These authors utilized the allelic combination *Su(H)*^1^/*Su(H)*^115B^, and observed a total block of CC differentiation accompanied by massive differentiation of PLs that invaded the MZ. According to the model proposed in the current study (**Fig. 8**), this is indeed the expected outcome of Tukusumi et al. experiments (Tokusumi et al., 2010), where a combined effect of Core Progenitor loss, along with inhibition of the CC fate and default differentiation towards the PL fate in Distal Progenitors is expected. It is still an open issue whether early fate specification in Distal Progenitors establishes the final (95:5) proportion of PLs versus CCs observed at the CZ, or if alternatively, the fate specification in Distal Progenitors brings about an initial 50:50 proportion of cells committed to one versus the other fate, followed by increased proliferative capacity of PL-committed Distal Progenitors.

The Notch pathway has been previously demonstrated to be necessary and sufficient for CC specification at the CZ (Duvic et al.; Lebestky et al., 2003; Mukherjee et al., 2011). Our results suggest that Notch-dependent CC specification occurs earlier, in Distal Progenitors, and that Notch is probably required again at later stages to sustain the original specification. Two works support this notion: First, Krzemien et al. (Krzemien et al., 2010b) utilized a clone labelling strategy at different larval stages, reaching the conclusion that blood cell progenitors of early L2 larvae, which have not yet developed a CZ, have already restrained their differentiating potential, either towards a PL or a CC fate. Second, at later stages of CC development, namely when Lz expression begins in unipotent CC progenitors at the CZ, it has been reported that Notch inhibits the expression of PL-specific genes (Terriente-Felix et al., 2013). In this study, gene expression manipulations performed with *lz*-Gal4 demonstrated that the Notch target gene *klumpfuss* (*klu*) mediates repression of P1 expression in CC progenitors, thereby stabilizing CC commitment (Terriente-Felix et al., 2013). We induced *klu* silencing with *domeMESO*-Gal4 in multipotent Distal Progenitors, and did not observe alterations of PL or CC differentiation (data not shown). This observation suggests that Notch target genes that regulate the early fate decision in Distal Progenitors, and those target genes that mediate CC-fate stabilization later at the CZ may be different. The identity of the Notch target genes that regulate initial cell fate determination in Distal Progenitors are not known and should be identified in future investigations.

Notch function as a regulator of the cell fate choice in Distal Progenitors is remarkably similar to the role of Notch reported in the control of a binary cell fate decision in mammalian lymphoid progenitors (Han et al., 2002; Liu et al., 2010; Pui et al., 1999; Radtke et al., 1999; Wilson et al., 2001). In this case, inhibition of the Notch pathway induces a default differentiation program towards a B lymphocyte fate, while the T lymphocyte fate is blocked (Han et al., 2002; Radtke et al., 1999; Wilson et al., 2001). Conversely, overactivation of the Notch pathway induces a T cell fate, while B cell differentiation is inhibited (Pui et al., 1999). Even though *Drosophila* progenitors are considered myeloid-like lineage cells, this similarity in Notch-dependent blood progenitor cell fate decision with the mammalian lymphoid lineage suggests a primitive mechanism of cell communication that determines a balance of blood cell types in hematopoiesis.

The Notch pathway is therefore employed several times during hematopoiesis at the lymph gland. First: A possible role in HSC at the 1^st^ larval instar (Dey et al., 2016); second: It is required for maintenance of Core Progenitors (this study); third: In Distal Progenitors it regulates the binary cell fate choice between PLs and CCs (this study); fourth: At the CZ, it is involved in fate stabilization in CC progenitors (Terriente-Felix et al., 2013), and later it is required for CC maturation and survival (Mukherjee et al., 2011). With the development of increasingly sophisticated genetic tools in mammalian systems, future studies may determine if Notch fulfills comparable context-specific functions throughout the mammalian hematopoietic hierarchy.

## Materials and Methods

### Fly Strains and crosses

The following Drosophila strains were used: *domeMESO*-GFP; *Hml*-dsRed; *domeMESO*-Gal4; *Antp*-Gal4; *UAS-NICD* (U. Banerjee), *Ser*-LacZ 9.5 (A. Bachmann); *eater*-dsRed; *eater*-Gal4 (Tsuyoshi Tokusumi); *hhF4*-GFP (RA. Schulz). The following stocks were obtained from the Bloomington Stock Center: *UAS-GFP* (#1521); *UAS-BFP* (#56807); *GFP RNAi* (#9331); *QUAS-GFP* (#52263); *QUAS-Gal80* (#51948); *Hml-Gal4* (#30139); *Hml-QF* (#66468); *Tub-Gal80*^*ts*^ (#7017); *UAS-Su(H)* (#17389); *UAS-Notch* (#26820); *UAS-Ser* (#5815), *Ubi-p63E(FRT.STOP)Stinger* (G-TRACE (#28280)(Evans et al., 2009); *UAS-rpr* (#50719); *UAS-Dcr2* (#24650 and #24651). *TepIV*-Gal4 was obtained from the Kyoto Stock Center (DGRC) (#105-442). All experimental crosses were performed at 25 °C and F1 larvae were incubated at 29°C to maximize Gal4 activity, with the exception of experiments involving *Tub-Gal80^ts^*, in which larvae were kept at 18 °C and then transferred to 29 °C to induce Gal4 activity. The PSC ablation protocol was performed at 25°C as previously described (Benmimoun et al., 2015).

The table below summarizes the RNAi lines utilized in this study, and whether or not they have been expressed together with a *UAS-Dicer2* construct (Dcr2). All RNAi constructs were expressed at 29ºC.

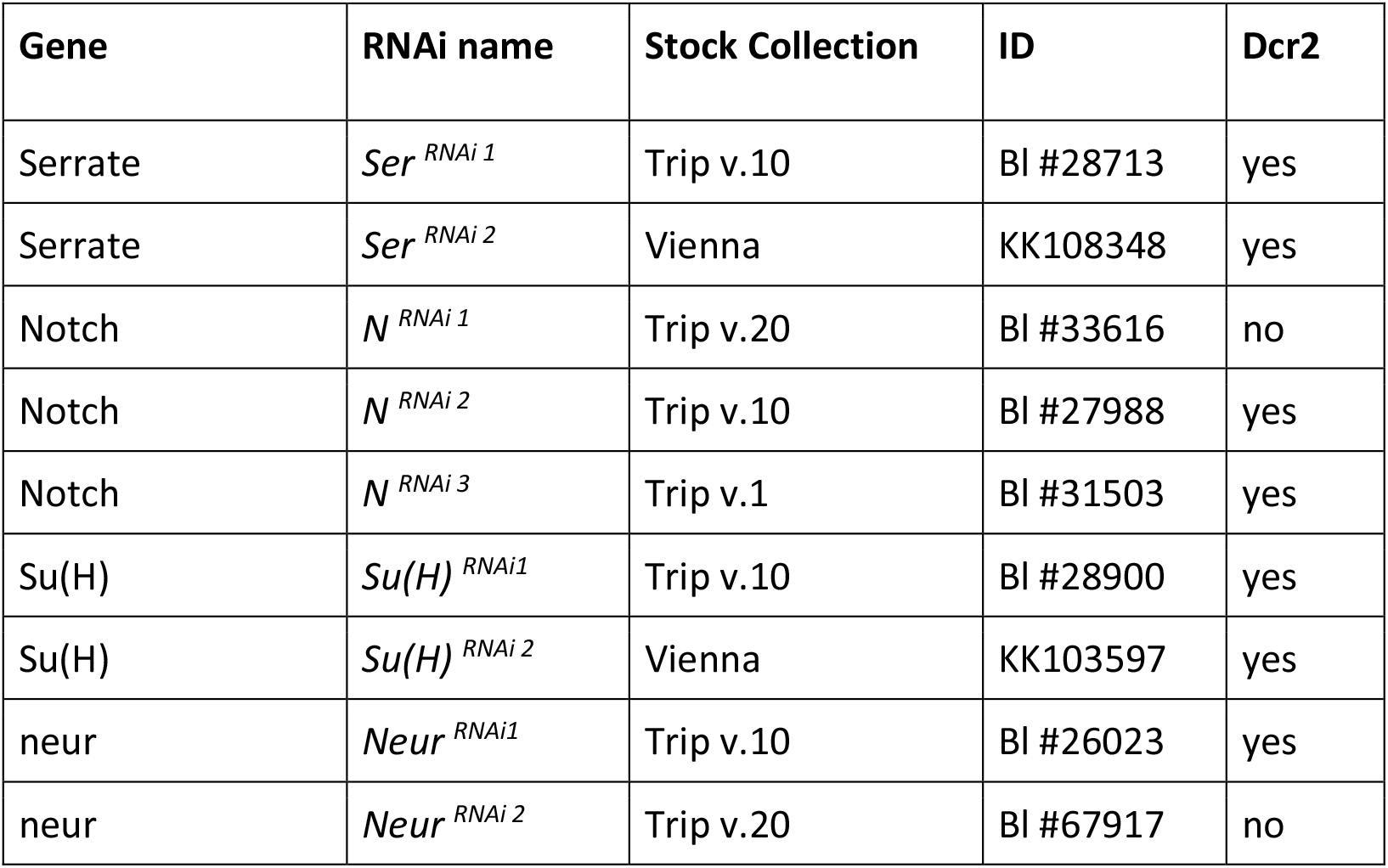

| Gene | RNAi name | Stock Collection | ID | Dcr2 |
| --- | --- | --- | --- | --- |
| Serrate | *Ser<sup>RNAi 1</sup>* | Trip v.10 | Bl #28713 | yes |
| Serrate | *Ser<sup>RNAi 2</sup>* | Vienna | KK108348 | yes |
| Notch | *N<sup>RNAi 1</sup>* | Trip v.20 | Bl #33616 | no |
| Notch | *N<sup>RNAi 2</sup>* | Trip v.10 | Bl #27988 | yes |
| Notch | *N<sup>RNAi 3</sup>* | Trip v.1 | Bl #31503 | yes |
| Su(H) | *Su(H)<sup>RNAi1</sup>* | Trip v.10 | Bl #28900 | yes |
| Su(H) | *Su(H)<sup>RNAi 2</sup>* | Vienna | KK103597 | yes |
| neur | *Neur<sup>RNAi1</sup>* | Trip v.10 | Bl #26023 | yes |
| neur | *Neur<sup>RNAi 2</sup>* | Trip v.20 | Bl #67917 | no |

### Immunofluorescence

Lymph glands were processed and stained as previously described (Evans et al., 2014): they were dissected from third-instar larvae (unless otherwise stated) in 1× PBS, fixed in 4% formaldehyde/1× PBS for 30 min, washed three times in 1×PBS with 0.4% Triton-X (1× PBST) for 15 min each, blocked in 10% normal goat serum/1× PBST for 30 min, followed by incubation with primary antibodies overnight at 4°C in blocking solution. Primary antibodies were washed three times in 1× PBST for 15 min each, re-blocked for 30 min, followed by incubation with secondary antibodies for 2 hr at room temperature. Samples were then washed three times in 1× PBST prior to mounting on glass slides in glycerol with Mowiol 4-88 anti-fade agent (EMD Millipore Corp., Billerica, MA). When required, lymph glands were stained with DAPI to visualize cell nuclei. To assess possible effects of genetic manipulations on cell size, the number of nuclei relative to the total *domeMESO* domain area of the lymph gland was quantified in single plane confocal images. The following primary antibodies and dilutions were used: rabbit α-βgal 1/5000 (Cappel), rabbit α-GFP 1/250 (ThermoFisher, Waltham, MA), mouse α-Lz 1/10, mouse α-Notch 1/10, mouse α-Antp 1/10 (Developmental Studies Hybridoma Bank, Iowa City, IA), Cleaved Caspase 3 1/100 (Cell signaling Technology), mouse αP1 1/100 (gift from I. Ando), rabbit α-ProPO 1/5000 (gift from G. Christophides), mouse α-Col 1/50 (gift of M. Crozatier). Alexa Fluor 488-, Alexa Fluor 647-, DyLight 405-and Cy3-conjugated secondary antibodies were used (Jackson Inmunoresearch, West Grove, PA). Reporters expressing GFP, BFP or dsRed were detected by fluorescence direct visualization.

### Image acquisition and processing

lymph glands were registered in a Zeiss LSM 510, 710 or 880 confocal microscope, either as Z-stacks or single confocal planes as indicated in each case. Images were processed using ImageJ software. Quantification of the area occupied by the indicated markers was performed using whole Z-projections of confocal stacks or single confocal planes as mentioned in each case. For single confocal planes, a stack from the central part of the lymph gland lobe was selected. Quantifications are expressed as the area occupied by the marker relative to the total lobe area. In the case of CCs, the total number of CCs relative to the total lobe area was quantified.

### Statistical analysis

Two-tailed unpaired Student’s t-tests or one-way ANOVAs (GraphPad Prism software) were used. The threshold for statistical significance was established as *p < 0.05, **p < 0.01 or ***p < 0.001. The “n” indicated in figure legends and throughout the paper refers to the number of lymph gland lobes analyzed in each experiment.

## Acknowledgements

We thank U. Banerjee lab members for sharing knowledge and expertise on lymph gland work. Schulz, A. Bachman and U. Banerjee for fly stocks; I. Ando, M. Crozatier and G. Christophides for the kind gift of α-P1, α-Collier and α-ProPO antibodies. The mouse α-z (U.Banerjee), mouse α-βgal (J.R. Sanes), mouse α-Notch (S. Artavanis-Tsakonas) and mouse α-Antp (D. Brower) antibodies, developed by the indicated investigators, were obtained from the Developmental Studies Hybridoma Bank, created by the NICHD of the NIH and maintained at The University of Iowa, Department of Biology, Iowa City, IA 52242. Stocks obtained from the Bloomington Drosophila Stock Center (NIH P40OD018537) and the Kyoto Stock Center were used in this study. We thank members of the Wappner lab for discussion and comments on the manuscript.

## Supplementary Figure legends

**Figure S1. Core progenitors localize in the internal region of the lymph gland.** Orthogonal projections of a confocal Z-stack image of primary lobes of the lymph gland of a wandering 3^rd^ instar larva are shown. Core progenitors are identified by the expression of BFP induced by *TepIV*-G4 (white; white-dashed line). Top panel: Composite image of *TepIV* > BFP (white; Core Progenitors), *domeMESO*-GFP (green; Progenitors) and *Hml*-dsRed (red; Cortical Zone). The red dotted line marks the outline of *Hml* expression. Middle panel: *TepIV* > BFP channel (Core Progenitors). Bottom: Schematics of the images; dark green: Core Progenitors; light green: Progenitors; red: Cortical Zone.

**Figure S2. Notch levels manipulation in Core Progenitors (A) Anti-Notch staining after *TepIV*-Gal4 driven silencing or overexpression of Notch.** Top: Notch (N) protein, as visualized with an anti-N antibody (αN; red) is homogeneously distributed throughout the lymph gland in control individuals that express GFP with a *TepIV*-Gal4 driver (green). Middle: lymph gland expressing a *N* RNAi driven by *TepIV*-Gal4 (*N^**RNAi 1**^*); N protein levels are reduced in the *TepIV* expression domain. Lower: *TepIV*-Gal4 driven overexpression of full-length N; N protein levels are increased in the *TepIV* expression domain. The dashed lines mark the outline of the two lymph gland lobes. Single-plane confocal images of lymph gland primary lobes of wandering 3^rd^ instar larvae are shown. Scale bars, 50 μm. **(B) Notch is required for Core Progenitor maintenance.** Experiment similar to the one depicted in Fig. 2B, where N expression was silenced with a different, non-overlapping, RNAi (*N*^***RNAi 2***^). *N*^***RNAi 2***^ expression in Core Progenitors, driven by *TepIV* Gal4, provoked loss of Core Progenitors, along with increased PLs and CCs. Quantification of whole Z-projection confocal images of primary lobes of wandering 3^rd^ instar larvae expressing *TepIV* > GFP and stained for P1 and ProPO (*p < 0.5, ***p < 0.001). Error bars represent SD. *TepIV* >, n = 11; *TepIV* > *N*^RNAi 2^, n = 10. **(C) Cell size is not affected by *Notch* silencing.** DAPI-stained nuclei lymph glands of either control or *domeMESO* > *N*^***RNAi 1***^ larvae were counted in the *domeMESO* > GFP^+^ area, and revealed no significant differences between the two groups. Single-plane confocal images of primary lobes from wandering third instar larvae were used. Error bars represent SD. (ns: Non-significant). *domeMESO* >, n=10, *domeMESO* > *N^RNAi 1^*, n = 10. **(D) *Notch* silencing in Core Progenitors does not affect the Posterior Signaling Center.** *TepIV*-Gal4 driven expression of a *Notch* RNAi (*N*^***RNAi 1***^) does not affect the Posterior Signaling Center, as revealed by an anti-Antennapedia (Antp) staining (red). *TepIV* > GFP expression (green) labels Core Progenitors. Upper: Control lymph gland without RNAi expression; Middle: Lymph gland that expresses *N*^***RNAi 1***^ driven by *TepIV*-Gal4. Single-plane confocal images of primary lobes from wandering third instar larvae are shown. Scale bars, 50 μm. Lower panel: Quantification of the number of Posterior Signaling Center cells per lobe (ns: Non-significant). Error bars represent SD. *TepIV* >, n = 8; *TepIV* > *N^RNAi 1^*, n = 8. (**E) Su(H) is required for Core Progenitor maintenance.** The experiment shown in Fig. 2C was repeated using a second non-overlapping RNAi. *Su(H)* ^RNAi 2^, whose expression was driven by *TepIV*-Gal4. As with *Su(H)* ^RNAi 1^, reduction of Core Progenitors and simultaneous increase of PLs was observed. Quantification of whole Zprojection confocal images of primary lobes from wandering third instar larvae expressing *TepIV* > GFP and stained for P1 and ProPO (*p < 0.05, ***p < 0.001). Error bars represent SD. *TepIV* >, n = 14; *TepIV* > *Su(H)*^RNAi2^, n = 19. **(F) Notch is not required for Core Progenitor maintenance at 1^st^ larval stage.** *N* silencing was induced in Core Progenitors with *TepIV*-G4 from mid L2 stage onwards using a *tub-Gal80^TS^* construct (*TepIV*^TS^). Top: Temperature shift protocol. Core Progenitors loss (green: *tepIV* > GFP), and enhanced differentiation of PLs (red: P1 staining) and CCs (cyan: ProPO staining) were observed. Upper: control lymph glands without RNAi expression; Middle: Lymph glands expressing *N^RNAi 1^*. Whole Z-projection confocal images of wandering third instar larvae are shown. Scale bars, 50 μm. Bottom: Quantification of the results (**p < 0.01). Error bars represent SD. *TepIV*^TS^>, n = 9; *TepIV^TS^* > *N*^RNAi 1^, n = 10.

**Figure S3. (A) Silencing of *Ser* at the PSC promotes differentiation of Crystal Cell and Plasmatocytes.** Experiment similar to the one depicted in Fig. 3A, utilizing second *Ser* non-overlapping RNAi (*Ser^RNAi 2^*), expressed under control of *Antp*>G4 driver. Whole Z-projection confocal images of primary lobes from wandering 3^rd^ instar larvae were quantified. PLs were visualized by anti-P1 staining and CCs by anti-ProPO staining. (***p < 0.001). Error bars represent SD. *Antp* >, n = 11; *Antp* > *Ser^RNAi 2^*, n = 11. **(B) Anti-Cleaved Caspase3 staining shows that cells expressing reaper are undergoing apoptosis.** The effect of expressing Reaper with an *Antp*-G4 driver was assessed by an anti-Cleaved Casp3 immunostaining. The Posterior Signaling Center (PSC) is marked with a yellow rectangle and zoomed below each photograph. *Antp*>: Control lymph gland without reaper expression; *Antp* > Rpr: Lymph glands expressing Reaper at the PSC. Note that apoptosis is triggered in the latter glands. **(C) Posterior Signaling Center of lymph glands of wild type individuals:** PSCs of 3 different glands (each one corresponding to a different row of pictures) are visualized with several markers simultaneously that include *Ser*-LacZ, *Antp* > GFP, *hh*-GFP, α-Col and α-Antp. Note that all the markers fully overlap with the exception of *Antp* > GFP, whose expressing is excluded from a subset of PSC cells. **(D) Apoptosis of Posterior Signaling Center cells after** *Col* **-induced expression of Reaper.** The PSC of lymph glands is marked with a yellow square and zoomed below each picture. *Col*>: Control lymph glands; *Col* > *rpr*: Lymph glands expressing Reaper under control of a *Col*-Gal4 driver. As can be seen by anti-cleaved caspase3 staining, Rpr expression provoked ablation of all the PSC cells, as revealed by the lack of cells that stain positive with an anti-Antp antibody. **(C) All the cells of the Posterior Signaling Center express***Col* **> GFP.** The PSC of a wild type lymph gland is marked with a yellow square and zoomed below. Anti-Antp staining reveals that *Col* > GFP is expressed in all PSC cells.

**Figure S4. (A) Notch silencing with a *domeMESO*-Gal4 driver utilizing an alternative *N* RNAi construct (*N^RNAi 3^*), also provoked increase of Plasmatocytes and reduction of Crystal Cells.** Expression of the *N^RNAi 3^* construct, with a different non-overlapping target sequence from that of *N^RNAi 1^*, was utilized to confirm the results of Fig. 5A. Quantification of whole Z-projection confocal images of primary lobes from wandering third instar larvae. PLs were visualized by anti-P1 staining and CCs by anti-ProPO staining. (*p < 0.05, ***p < 0.001). Error bars represent SD. *domeMESO* >, n = 8; *domeMESO* > *N^RNAi 3^*, n = 5. **(B) *Su(H)* silencing with the *domeMESO*-Gal4 driver, utilizing an alternative RNAi construct (*Su(H)^RNAi 2^*) recapitulates the effect on differentiation of Plasmatocytes and Crystal Cells.** The results shown in Fig. 5B were confirmed by using the *Su(H)*^RNAi 2^ construct with a different non-overlapping target sequence from that of *Su(H)*^RNAi 1^. Quantification of whole Z-projection confocal images of primary lobes from wandering third instar larvae. PLs were visualized by anti-P1 staining and CCs by anti-ProPO staining. (**p < 0.01, ***p < 0.001). Error bars represent SD. *domeMESO* >, n = 12; *domeMESO* > *Su(H)*^RNAi 2^, n = 12. **(C) Notch over-expression in *domeMESO*-positive progenitors.** Anti-Notch (N) staining (red) was utilized to show an increase of Notch levels in progenitors (visualized in green; *domeMESO* > GFP) following *domeMESO*-G4 driven over-expression of full-length Notch. Upper: Control lymph glands; Lower: lymph glands that over-express N. Single-plane confocal images of primary lobes from wandering third instar larvae are shown. Scale bars, 80 μm. **(D) *Notch* silencing in *domeMESO* positive progenitors which do not express** *Hemolectin* (*Hml*) show increase of Plasmatocytes and reduction of Crystal Cells. *N^RNAi 1^* was expressed with a *domeMESO*-Gal4 driver, together or not with the Gal4 inhibitor Gal80 in *Hml*-positive cells (utilizing *Hml*-F driven expression of *QUAS-Gal80*). PLs are visualized in red (P1 staining) and CCs in cyan (ProPO staining). Upper: control lymph glands; 2^nd^ row: lymph glands expressing *N^RNAi 1^*, without Gal80 expression; 3^rd^ row: lymph glands expressing *N^RNAi^* ^1^ along with Gal80 in *Hml*-positive cells. Whole Z-projection confocal images of primary lobes from wandering third instar larvae are shown. Scale bar, 80 μm. Lower panels: Quantification of the indicated markers. The letters indicate statistical difference (Tukey′s multiple comparison test). Error bars represent SD. *domeMESO*-G4 >, *Hml*-QF >, n = 10; *domeMESO*-G4 > *N^RNAi 1^*, *hml*-QF >, n = 10; *domeMESO*G4 > *N^RNAi 1^*, *Hml*-QF > Gal80, n = 14. **(E) *Hml-* QF driven expression of *QUAS-Gal80* is effective in repressing Gal4 activity at the Cortical Zone.** Gal80 is effective in inhibiting Gal4 activity, as assessed by a virtual block of *Hml*-Gal4 driven *UAS-GFP* expression (green) when Gal80 is expressed in the same cells using *Hml*-QF > *QUAS-Gal80* (lower panel). Upper panel: control lymph glands without expression of Gal80. DAPI staining shows nuclei. Single-plane confocal images of wandering third instar primary lymph gland lobes are shown. Scale bars, 80 μm.

**Figure S5. Notch does not affect Plasmatocyte or Crystal Cell differentiation after *eater* expression has started.** Pictures show expression of the PL marker P1 (red) and the CC marker ProPO (cyan) in control lymph glands (upper panels) and in lymph glands with *eater*-Gal4 driven expression of *Notch* RNAi (*N^RNAi 1^*)(2^nd^ row panels) or of full-length Notch (3^rd^ row panels). Whole Z-projection confocal images of primary lobes from wandering third instar larvae are depicted. Scale bar, 60 μm. Lower panels: Quantification of the indicated markers. Same letter indicates no statistical difference between means (Tukey′s multiple comparison test). *eater* >, n = 8; *eater* > N^RNAi 1^, n = 10; *Eater* > N, n = 10.

**Figure S6. (A) Silencing of *Ser* at the Medullary Zone with an alternative RNAi also impairs Crystal Cell differentiation and enhances the Plasmatocyte fate.** A second non-overlapping RNAi targeting *Ser* (*Ser*^*RNAi* 2^) was utilized to confirm the results depicted in Fig. 7C. Whole Z-projection confocal images of primary lobes from wandering third instar larvae were quantified. PLs were visualized through P1 staining and CCs through ProPO staining. (***p < 0.001). Error bars represent SD. *domeMESO* >, n = 12; *domeMESO* > *Ser*^RNAi 2^, n = 10. **(B) A second RNAi targeting***neutralized*, **expressed at the Medullary Zone, also decreases differentiation of Crystal Cells while increasing Plasmatocytes.** The results shown in Fig. 7D were validated by utilizing an alternative RNAi targeting a different non-overlapping sequence of the gene *neur* (*neur^RNAi 2^*), under the control of *domeMESO*-Gal4. Whole Z-projection confocal images of primary lobes from wandering third instar larvae were quantified. PLs were visualized through P1 staining and CCs through ProPO staining. (**p < 0.05, ***p < 0.001). Error bars represent SD. *domeMESO* >, n = 10; *domeMESO* > *neur*^RNAi 2^, n = 14.

