## Supplementary Figures for "Context-specific functions of Notch in *Drosophila* blood cell progenitors"

### Sup Figure 1
(1 column)

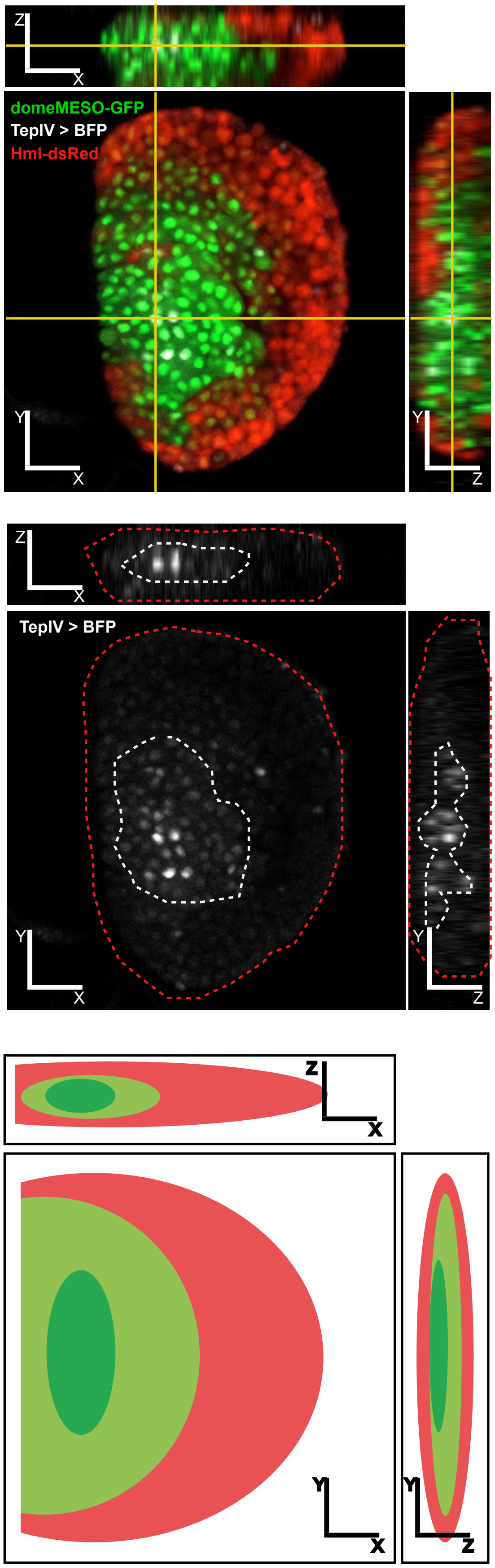

### Sup Figure 2
(2 columns)

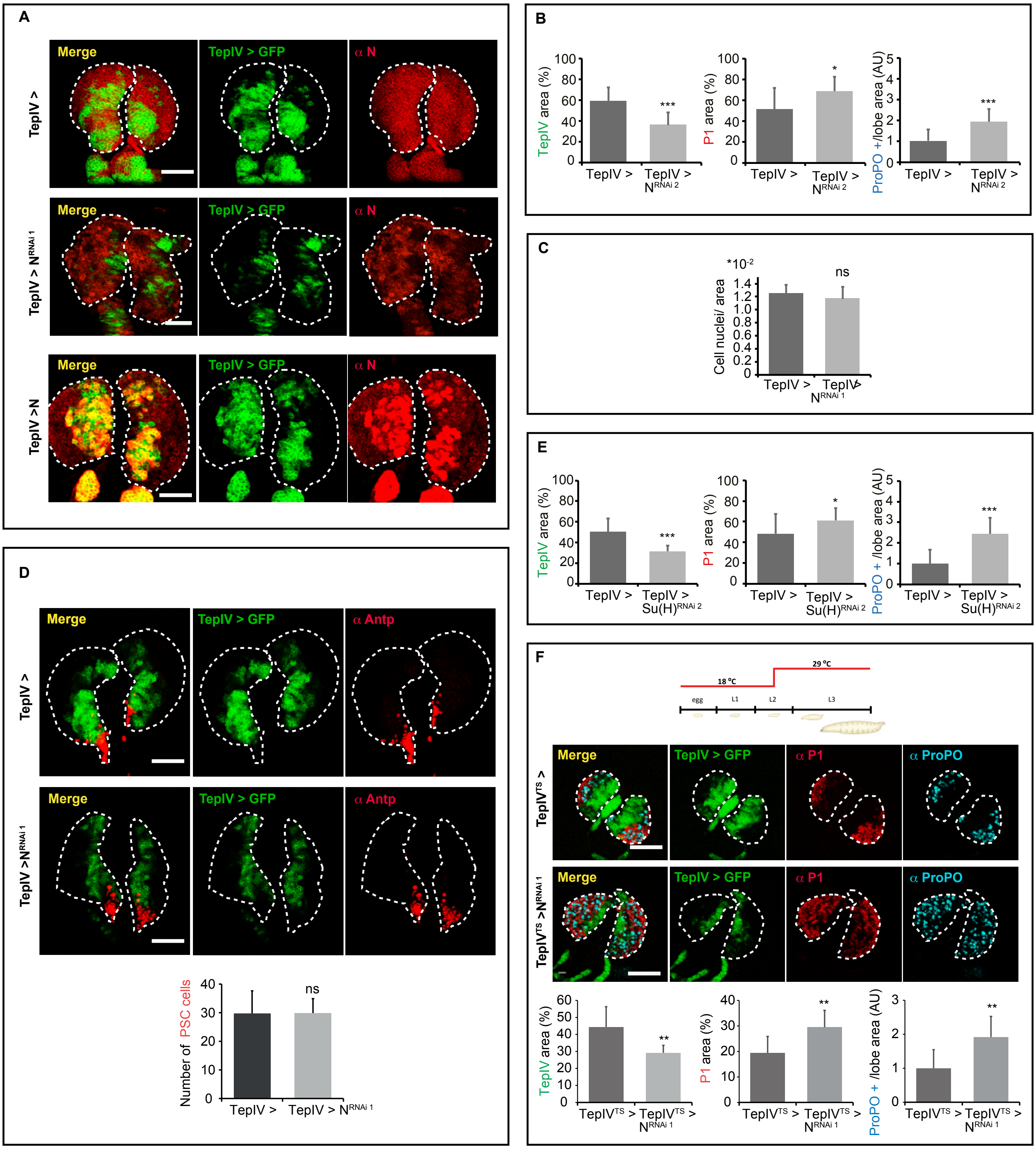

### Sup Figure 3
(2 columns)

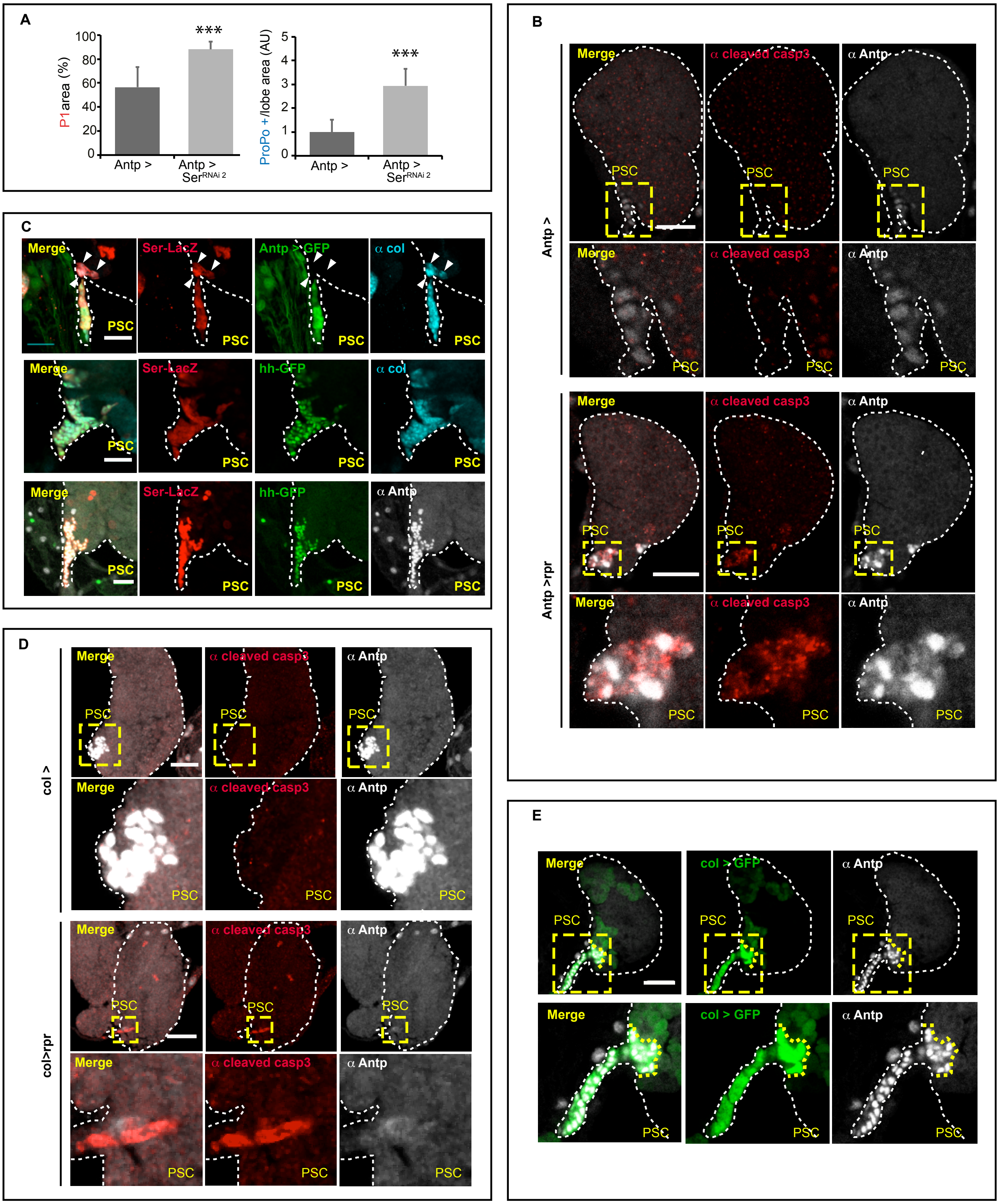

### Sup Figure 4
(2 columns)

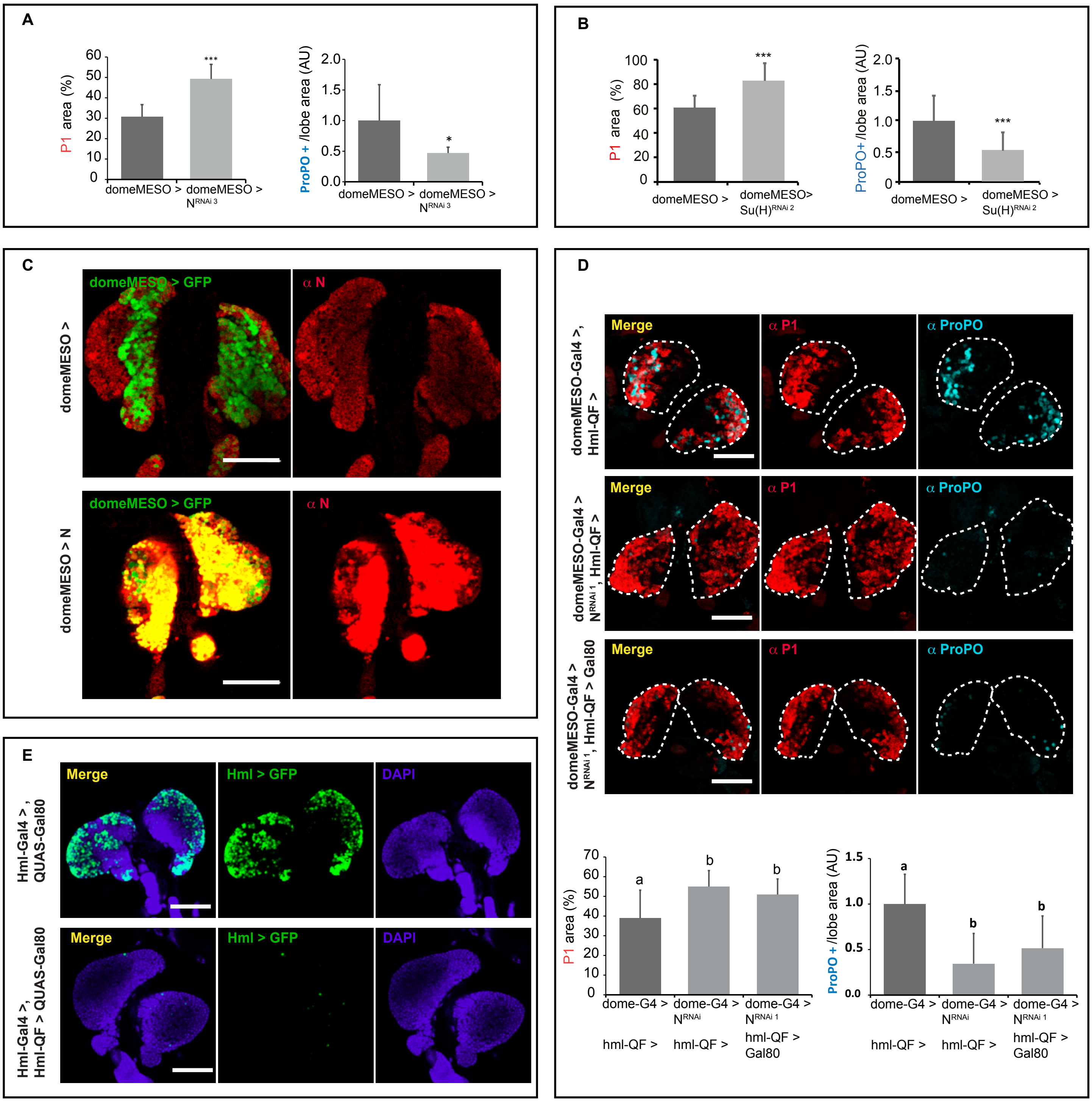

### Sup Figure 5
(1 column)

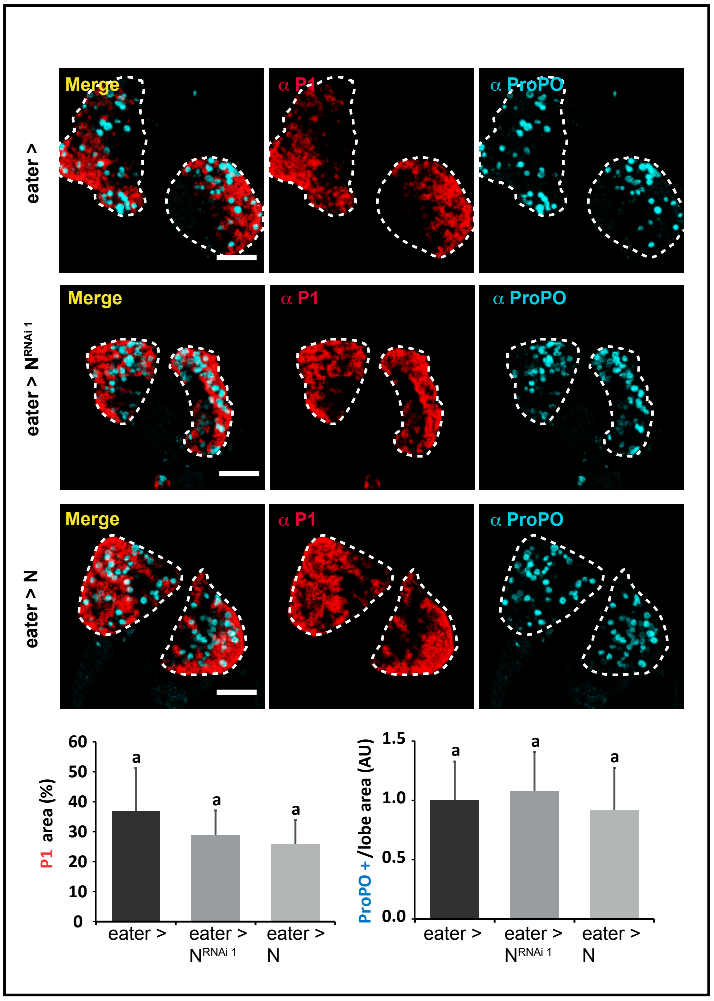

### Sup Figure 6
(1 column)

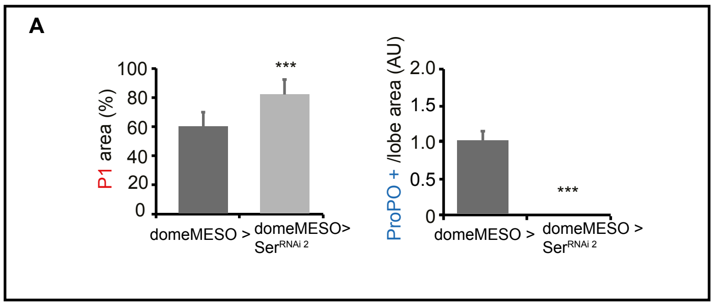


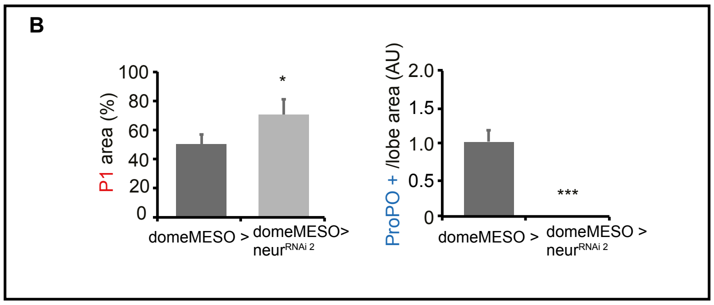
